# Learning Perturbation Effects Through Contrastive Alignment of Multimodal Biological Embeddings

**DOI:** 10.64898/2026.06.23.734145

**Authors:** Wenxin Long, Tianyu Liu, Artur Szałata, Fabian J. Theis, Lingzhou Xue, Hongyu Zhao

## Abstract

Multimodal single-cell perturbation screens offer a scalable approach for characterizing the effects of genetic and chemical interventions on cellular state. However, most existing representation-learning methods are tailored to a single perturbation modality and fail to explicitly incorporate external semantic knowledge, which limits their ability to generalize across datasets and perturbation types. Here, we introduce PertOmni, a CLIP-style multimodal representation-learning framework that aligns transcriptomic perturbation signatures with text-derived embeddings of curated genes and compound descriptions, as well as image-derived embeddings from cell paintings. PertOmni jointly trains a shared transcriptomic encoder and dataset-specific text encoders using a masked contrastive objective that emphasizes within–cell-type discrimination while mitigating confounding effects arising from cell-type heterogeneity. We evaluate the produced joint embedding space on bi-directional retrieval, drug–gene interaction inference, and perturbation prediction across both small-molecule and CRISPRi perturbation datasets, and demonstrate consistent improvements over strong baseline methods.

## 1. Introduction

Numerous perturbation experiments have been developed to systematically quantify cellular changes under varying conditions (1–4), enabling the study of how cells respond to interventions. Recent advances in single-cell RNA sequencing enable these responses to be profiled at single-cell resolution, yielding control and perturbed transcriptomic readouts across many perturbations (2, 5, 6). Broadly speaking, such perturbations can be grouped into three major categories based on the perturbation used. Genetic perturbations, such as CRISPR-based ones (2), leverage targeted genome editing to disrupt or modulate gene function, thereby altering the expression of the perturbed gene and its associated regulatory network. Small-molecule perturbations (7, 8), in contrast, use chemical compounds that typically act on protein products such as enzymes or receptors, indirectly reshaping the transcriptional landscape of affected pathways. Biologics form another category and include, for example, extracellular ligand stimulation with cytokines. Together, these experimental strategies have become important tools for understanding cellular circuitry, providing insights into fundamental biology and enabling the discovery of therapeutic targets.

Analyzing gene expression profiles from perturbation experiments remains challenging. Historically, the high cost of such experiments limited the availability of large-scale, genome-wide, or atlas-level datasets (6). Although several institutes and companies have released single-cell perturbation datasets with multiple replicates and experimental conditions (8–11), data quality and noise remain persistent concerns. Perturbations can trigger stress or cell-death programs; moreover, nuclease-based genetic perturbations can induce complex DNA damage such as chromothripsis (12), and perturbation efficacy can be heterogeneous across cells (13). In addition, single-cell RNA-seq measurements are prone to technical artifacts and batch effects (14, 15). These challenges present as obstacles in analyzing perturbation effects rigorously.

In addition to transcriptomic profiling, image-based perturbation assays such as Cell Painting provide a complementary strategy for measuring cellular responses (16). Cell Painting uses multiplexed fluorescence microscopy to capture high-content morphological profiles across multiple cellular compartments, enabling large-scale phenotypic screening of genetic or chemical perturbations (17, 18). Compared with single-cell RNA-seq, Cell Painting offers a different view of perturbation effects by directly measuring cellular morphology and subcellular organization. However, linking image-derived phenotypes with textual biological knowledge and transcriptomic perturbation effects remains underexplored, especially in settings involving both genetic and small-molecule perturbations.

A range of computational methods for denoising, normalizing, and modeling perturbation responses has been proposed to address these challenges, including (19–26). These methods typically use as input a representation of the perturbation and the transcriptomic profile of the control cells of interest. More recently, several single-cell foundation models such as (27–30) have also claimed to support perturbation response prediction, but benchmarking studies such as (10, 31, 32) suggest that their generalization remains limited. scGen (33) aims to predict single-cell perturbation responses by learning a latent representation and transferring perturbation effects via latent-space arithmetic, while GPerturb (34) estimates gene-level perturbation effects with calibrated uncertainty. Despite these advances, two major gaps remain. First, most approaches overlook the integration of external biological knowledge expressed in natural language, such as functional annotations of genes and drugs, which could improve interpretability and predictive accuracy. Second, few frameworks model both genetic and chemical perturbations. Such integration would enable the inference of drug–target relationships.

Therefore, we introduce PertOmni, a unified framework for modeling and bridging perturbation data across diverse perturbations and technologies. PertOmni learns a shared embedding space by contrastively aligning LLM-derived perturbation text embeddings with transcriptomic perturbation-effect embeddings across genetic and small-molecule screens. Specifically, we embed curated gene and drug descriptions, such as from NCBI (35), and jointly train text and transcriptome encoders so that matched perturbations and expression readout are close in the latent space. We evaluate PertOmni on (i) bi-directional perturbation retrieval (text to transcriptome and transcriptome to text) and (ii) downstream applications, including drug–gene interaction inference and retrieval-based perturbation prediction, demonstrating improved alignment and utility compared to strong baselines. Codes of PertOmni can be found here ^1^.

## 2. Results

### Method overview

Figure 1 summarizes PertOmni, which learns a joint embedding space aligning (i) perturbed transcriptomic profiles and (ii) perturbation identities across genetic and small-molecule screens. PertOmni uses a shared transcriptome encoder and perturbation-type–specific text encoders (gene vs compound) that map LLM-initialized textual descriptions into the same latent space. Training uses a masked contrastive objective that forms negatives only within the same cell line, reducing confounding from cross–cell-line heterogeneity while preserving fine-grained within-context discrimination.

**Figure 1:**
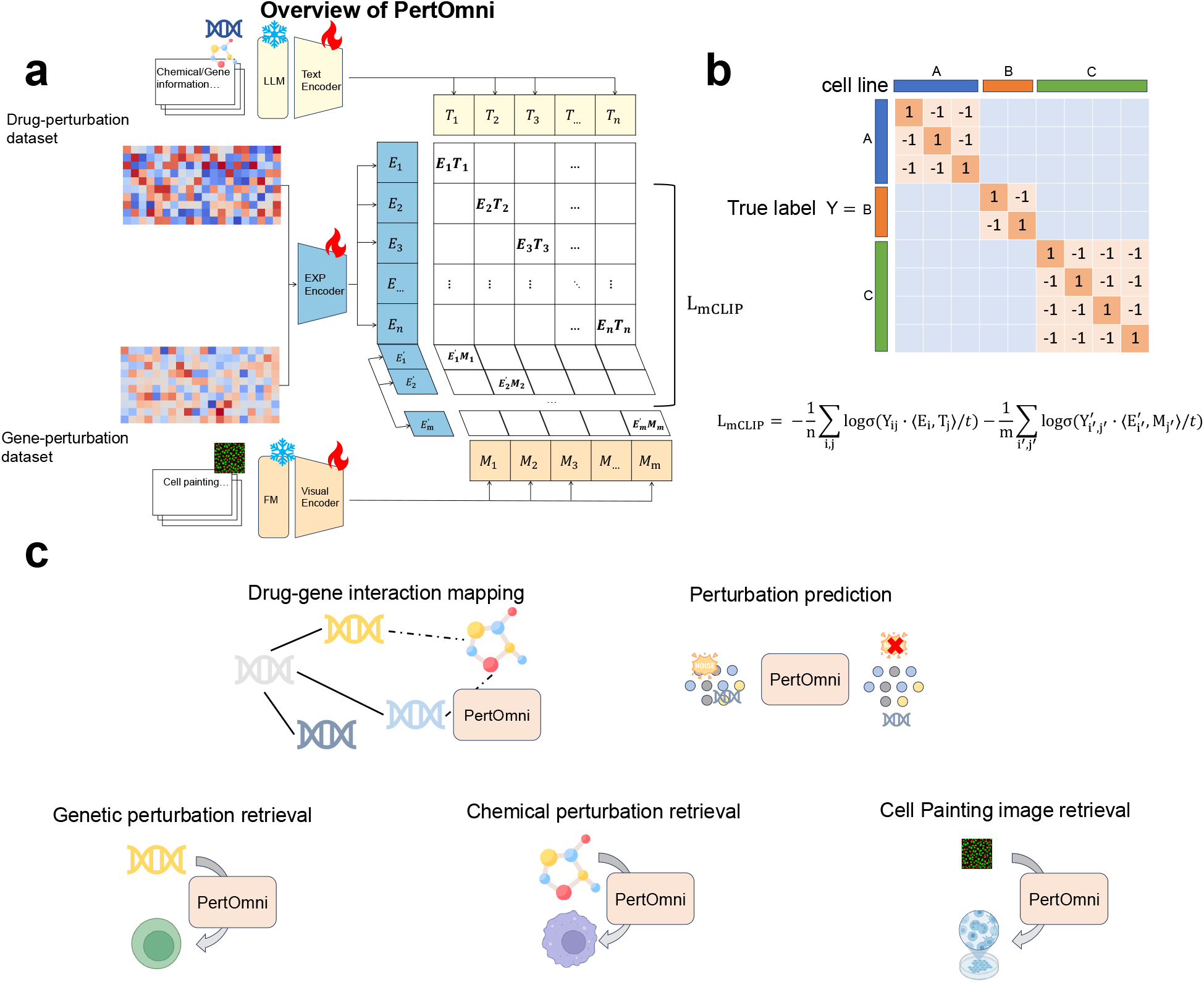
Overview of PertOmni. (a) The illustration of the model architecture. We leverage contrastive learning and modality-specific encoders to build PertOmni. (b) The illustration of masked contrastive learning. We modify the CLIP loss, *L*_mCLIP_, to leverage multimodal and masked information during training. (c) Task illustration of downstream applications.

We evaluate the learned embeddings on four downstream applications: (i) drug–gene interaction inference (36) via fine-tuning the text encoders on DGIdb labels (Figure 2), (ii) retrieval-based perturbation effect prediction using nearest neighbors in the learned space (Figure 3), (iii) bi-directional cross-modality retrieval between text and transcriptome embeddings (Figure 4), and (iv) bi-directional cross-modality retrieval between text and image embeddings (Figure 5). Additional ablations on transcriptome embedding choices are shown in Supplementary Figure 1.

**Figure 2:**
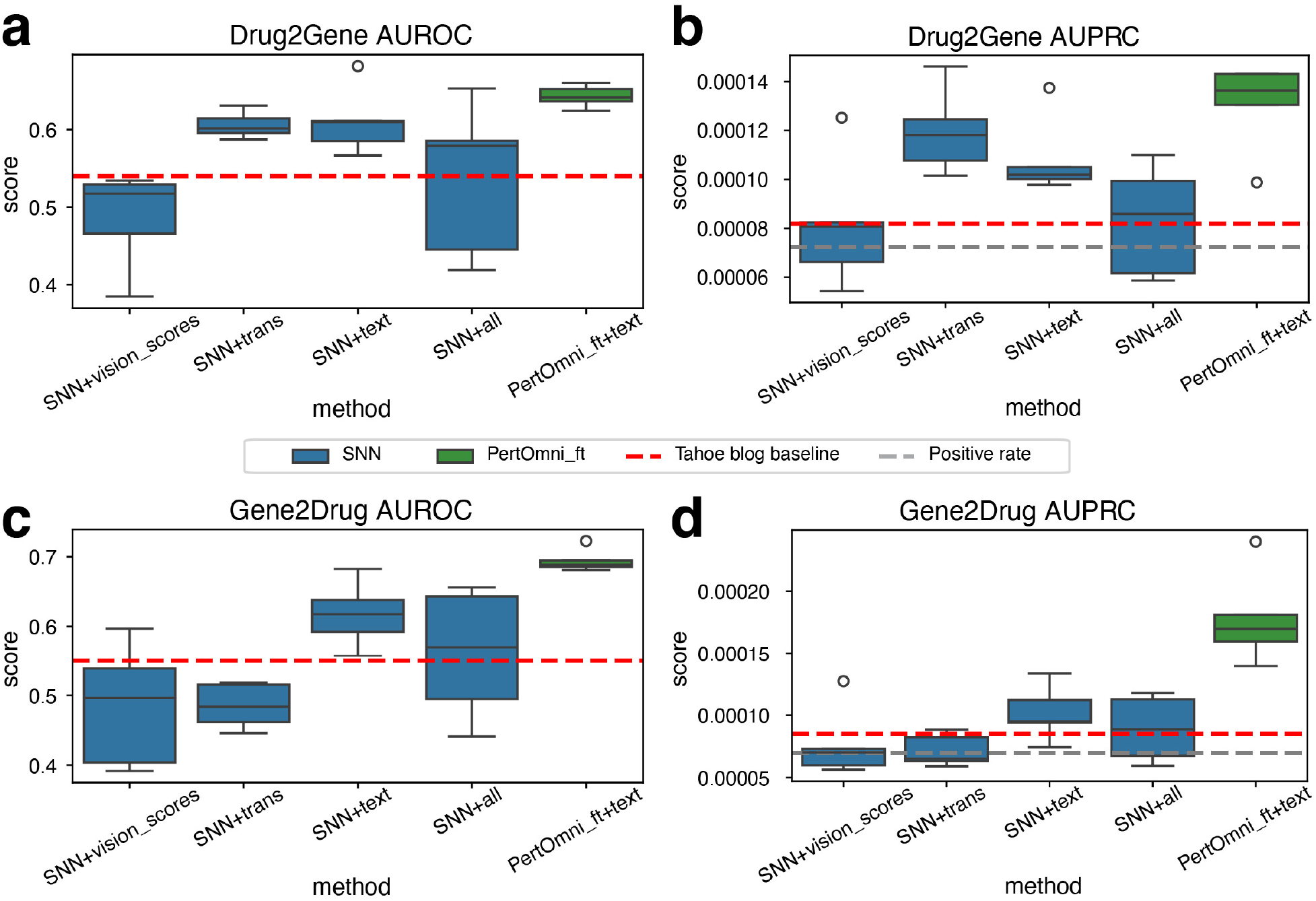
Results of drug-gene pair prediction. Details of choices are summarized in the legend. Tahoe blog baseline first computes differential Vision Scores relative to control cells, and then predicts drug-gene matching using cosine similarity between differential Vision Scores of drugs and genes. (a) AUROC comparison for the Drug2Gene task. (b) AUPRC comparison for the Drug2Gene task. (c) AUROC comparison for the Gene2Drug task. (d) AUPRC comparison for the Gene2Drug task.

**Figure 3:**
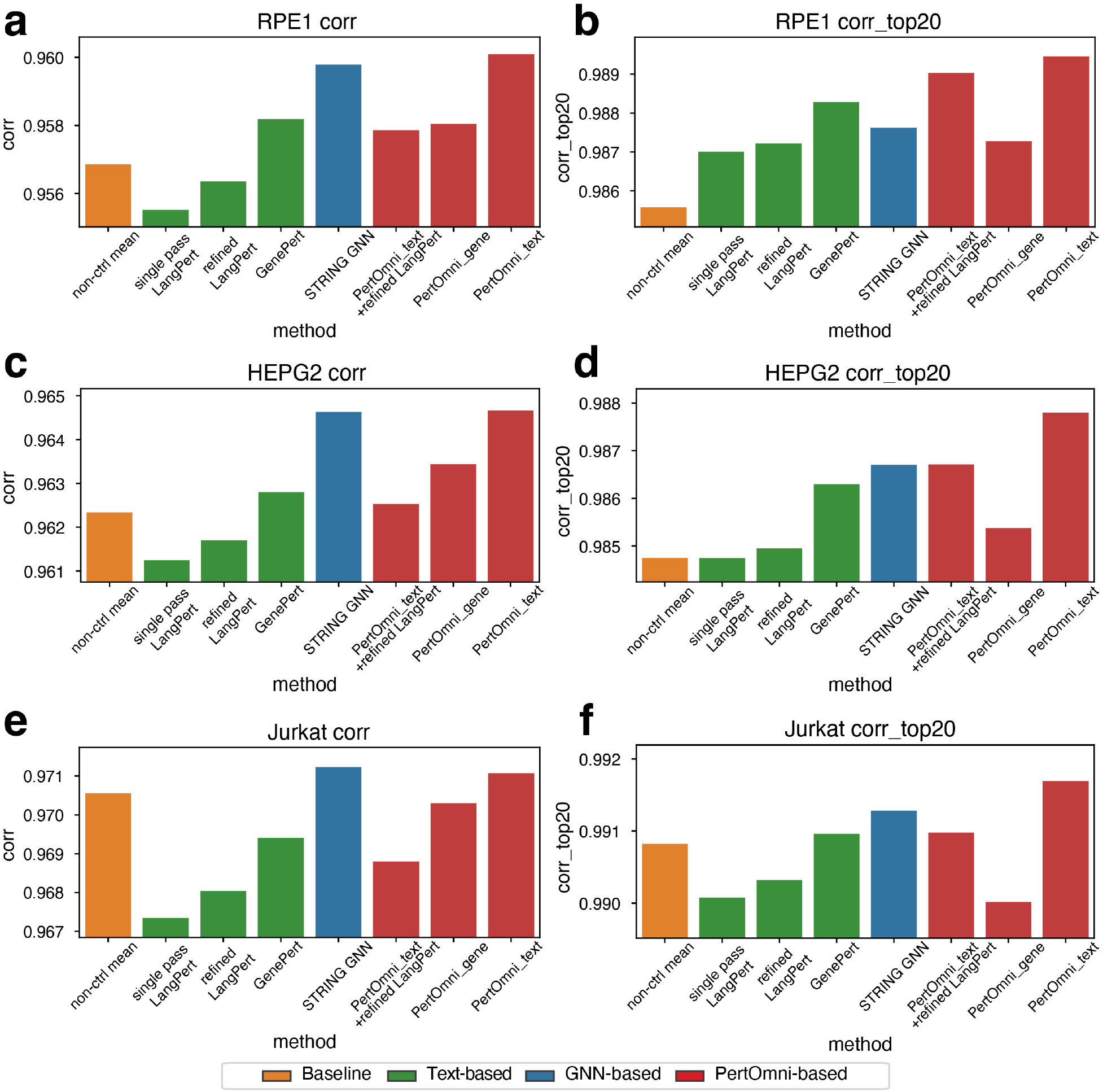
Perturbation prediction via retrieval. Details of methods are summarized in the legend. Performance is measured as the Pearson correlation between predicted and observed gene expression. (a) All genes, RPE1. (b) Top-20 DEGs, RPE1. (c) All genes, HepG2. (d) Top-20 DEGs, HepG2. (e) All genes, Jurkat. (f) Top-20 DEGs, Jurkat.

**Figure 4:**
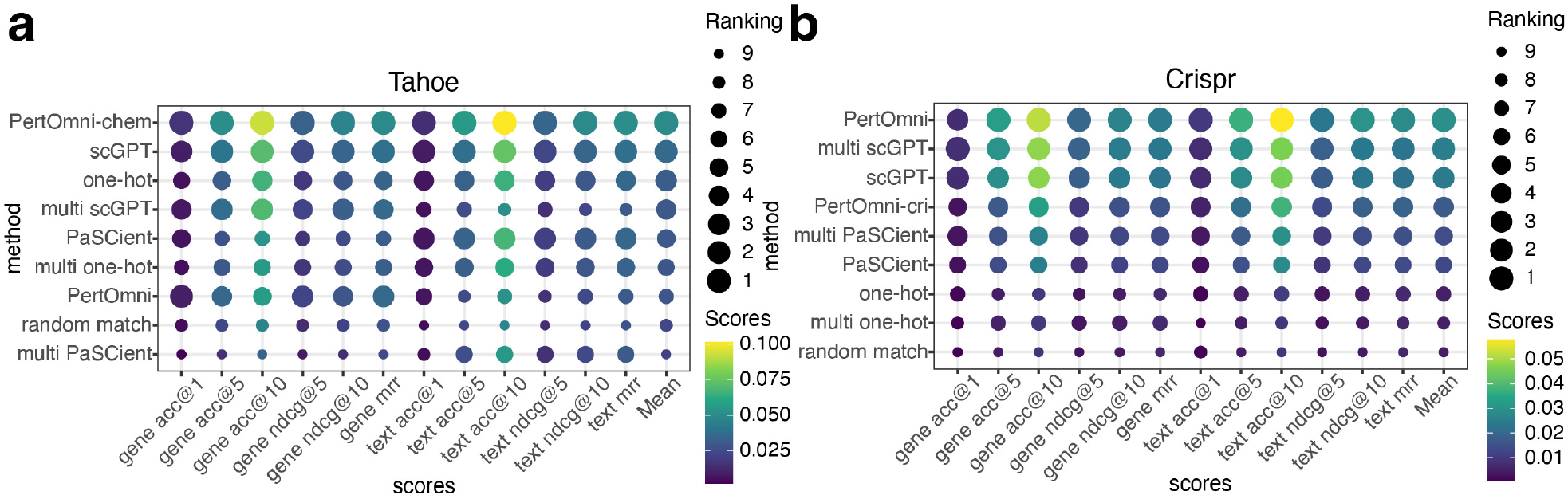
Results of cross-modality retrieval between transcriptomic and text. We find that using training data from two perturbations can benefit CRISPR-based retrieval, whereas it does not improve Chem-based retrieval (Tahoe). (a) Multi-metric comparison based on the Tahoe 100M dataset. (b) Multi-metric comparison based on the merged CRISPR dataset.

**Figure 5:**
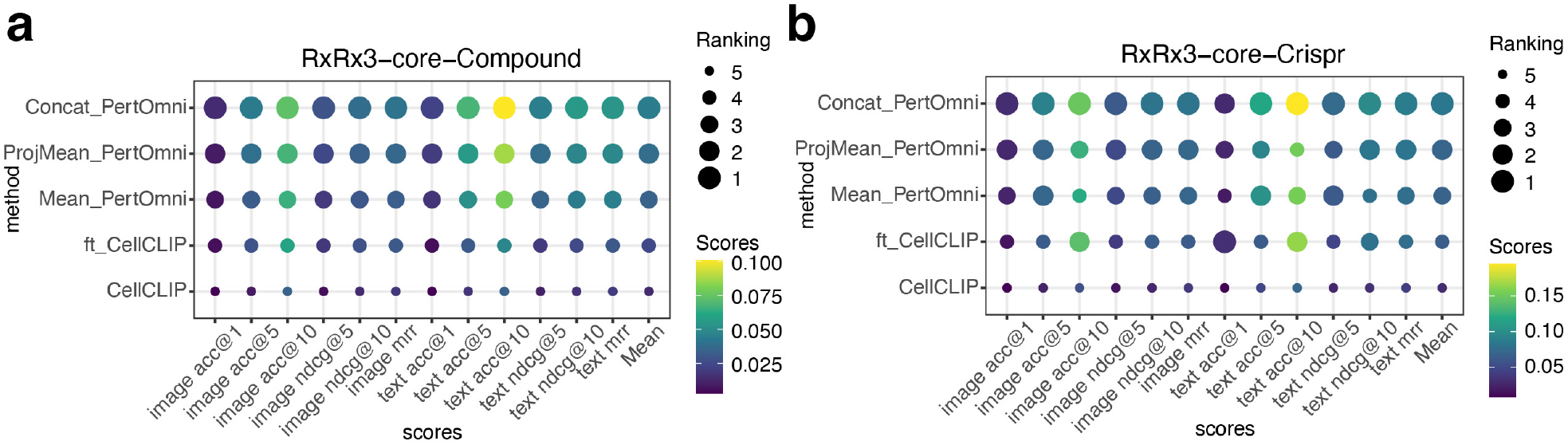
Results of cross-modality retrieval between image and text. (a) Multi-metric comparison based on the compound perturbations in RxRx3-core dataset. (b) Multi-metric comparison based on the genetic perturbations.

### Predicting drug–gene interaction with PertOmni

We fine-tune PertOmni for drug–gene interaction inference by optimizing a retrieval-style objective that increases similarity for annotated DGIdb drug–gene pairs while leaving unannotated pairs untreated (positive–unlabeled setting). We split drugs and genes so that test drugs/genes are unseen during training, and evaluate two settings: Drug2Gene (rank genes for a drug) and Gene2Drug (rank drugs for a gene). Baselines include similarity-based matching (37), and an SNN classifier trained on Vision Scores (38) or PertOmni text, transcriptome, or text+transcriptome embeddings.

As shown in Figure 2(a–d), *PertOmni_ft+text* achieves the best performance across both tasks, improving AUROC and AUPRC over all similarity-based baselines and over the SNN classifier. Supplementary Figure 2 shows that fine-tuning substantially improves over the zero-shot variant, indicating that aligning embeddings with interaction supervision is important beyond generic retrieval alignment.

Because DGIdb annotations are sparse, we report AUPRC alongside the task positive rate and interpret gains relative to this baseline. Drug2Gene is consistently harder than Gene2Drug, consistent with the many-to-many nature of drug targeting and incomplete interaction annotations: a single compound can have multiple true targets, and many true targets may be unlabeled.

### Predicting perturbation effects with PertOmni

We evaluate retrieval-based perturbation prediction by querying the learned embedding space for nearest-neighbor perturbations and aggregating their observed expression profiles to predict the held-out perturbation response. Unlike LLM-only approaches such as LangPert (39), PertOmni provides an explicit, data-grounded neighborhood over perturbations that can be used directly for kNN prediction and for defining gene sets.

Figure 3(a–f) reports Pearson correlation between predicted and observed expression across three perturb-seq datasets (RPE1 (9), HepG2, Jurkat (40)), evaluated on either all genes or the top-20 DEGs. Across settings, kNN prediction using PertOmni text embeddings yields the best performance with *k* = 100. The comparison of PertOmni text embeddings with different *k*s can be found in Supplementary Figure 3. From the linecharts, the performance is sensitive to the neighborhood size *k*. At larger *k*, transcriptome-derived embeddings become competitive with text embeddings, suggesting that denser neighborhoods partially compensate for weaker semantic priors. PertOmni consistently outperforms LangPert when evaluated on sufficiently large gene sets; moreover, LangPert improves when restricted to gene sets proposed by PertOmni (PertOmni+refined LangPert), indicating that the gene-selection signal learned by PertOmni is useful beyond our framework. We include the non-control (non-ctrl) mean as a strong baseline. While competitive at small *k*, this baseline is consistently surpassed by PertOmni as *k* increases, whereas other methods, including LangPert, do not reliably outperform it.

Additionally, we compared our proposed method with GenePert (41) and STRING GNN (42). GenePert (41) fits a ridge regression model from GenePT embeddings of perturbed genes (43) to mean expression profiles and then uses the learned mapping to predict expression profiles for unseen perturbations. STRING GNN, one of the top-performing methods in the benchmark studies from (42), initializes a GNN with weights pretrained by link prediction on STRING (44), extracts the embedding of each perturbed gene from the pretrained model, and uses a linear head to predict the observed treatment-effect vector by minimizing squared error. From Figure 3, our proposed PertOmni_text outperforms all other methods.

### Cross-modality retrieval benchmarking on transcriptomic and text

We assess how well PertOmni aligns transcriptomic and textual representations using bi-directional retrieval (text ↔ transcriptome) under two perturbation regimes: chemical-only training on Tahoe-100M (8), CRISPRi-only training on merged perturb-seq datasets (9, 40, 45), and joint training on both. We compare against single-modality baselines (e.g., scGPT (27), PaSCient (46)), one-hot embeddings, and random matching. Retrieval is evaluated with ACC@K, nDCG@K, MRR, and an aggregate mean score.

Figure 4 summarizes performance with a bubble plot across metrics. PertOmni and its variants achieve the top or near-top rank on most metrics in both Tahoe and CRISPR settings, indicating robust cross-modal alignment rather than gains limited to a single modality. Detailed results are in Supplementary Figures 4 and 5.

Joint training improves CRISPR-side retrieval while slightly degrading Tahoe-side retrieval. We hypothesize that this reflects data imbalance, as we have substantially more CRISPR perturbations, which can bias the shared space toward the higher-sample data. Exploring explicit reweighting or balanced batching is a natural next step. Architecture and hyperparameter ablations are reported in Supplementary Figures 1, 6, 7, 8, 9, 10, 11, 12, 13, and 14.

### Cross-modality retrieval benchmarking on image and text

We further assess PertOmni’s ability in aligning image and textual representations. We compare PertOmni with CellCLIP (47) on RxRx3-core (48), a Cell Painting dataset curated from the RxRx3 dataset (18). Following the preprocessing steps in CellCLIP, we use only 5 channels in this dataset, and get channel-wise 1536-dimensional embeddings using pretrained DINOv2 (49).

On the image side, we train three different channel aggregation methods for PertOmni: (i) Concat_PertOmni: concatenates the channel-wise embeddings into a 5 × 1536 dimensional representation; (ii) ProjMean_PertOmni: projects each channel embedding with a shared MLP, and then mean-pools the projected channel embeddings. (iii) Mean_PertOmni: direct averages across channels. Then we average the image embeddings to perturbation-level, and optimize the contrastive loss between the perturbation-level image embedding and the LLM-derived text embedding. For comparison with CellCLIP, we include both the pretrained checkpoint provided by CellCLIP and CellCLIP fine-tuned on the RxRx3-core dataset (ft_CellCLIP). Retrieval between image and text is evaluated at the perturbation level separately for compound and CRISPR perturbations, with ACC@K, nDCG@K, MRR, and an aggregate mean score.

Figure 5 summarizes performance with a bubble plot across metrics. All three variants of PertOmni perform better than CellCLIP and ft_CellCLIP in the aggregated mean score in both compound and genetic perturbations, indicating its robustness of aligning between image and text modalities. Among the three variants, Concat_PertOmni performs the best, followed by ProjMean_PertOmni and then Mean_PertOmni. This suggests that preserving channel-specific information is important for cross-modality alignment. Concat_PertOmni retains the full channel identity by concatenating channel-wise embeddings, whereas ProjMean_PertOmni preserves channel-wise processing but compresses the channels through mean pooling. Mean_PertOmni performs the strongest aggregation by directly averaging across channels, which removes channel identity earliest and may obscure stain-specific morphological signals. Detailed results can be found in Supplementary Figures 15, 16.

## 3. Discussion

We introduce PertOmni, a framework for aligning representations of perturbation effects across modalities. By leveraging LLM embeddings and masked contrastive learning, PertOmni successfully integrates heterogeneous cell context and perturbation information, outperforming strong baselines across four applications relevant to drug discovery.

The key innovation of PertOmni lies in two dimensions. First, PertOmni leverages the prior knowledge introduced by databases and translates the textual descriptions into numerical representations with the help of LLM embedding layers. We also perform a joint training approach by matching the samples from sequencing data with their correct paired embeddings in different cell types, and thus we can achieve perturbation modeling at the cell-type-level resolution, thereby better aligning with biological requirements and providing more insights. Second, our approach attempts to integrate perturbation datasets from diverse sources, including genetic perturbations and chemical perturbations, thereby directly linking drugs to their potential targets. We also demonstrated that after integrated training, we can better predict drug-gene interactions, thereby providing insights for clinical applications.

In addition, PertOmni simultaneously addresses multiple applications, thereby supporting its role as a unified model for studying perturbation effects. In addition to predicting drug-gene interactions, we can also enhance the correlation of predicting perturbations effects at the cellular type level under different gene groups. PertOmni also enables us to search for neighbors of specified genes based on embedding similarity, thereby enhancing the effectiveness of related tools by constructing gene similarity networks. For example, utilizing improved gene sets can help LangPert better predict perturbation outcomes. Additionally, PertOmni shows its superiority in multi-modality tasks, outperforming baselines in both gene-text retrieval and image-text retrieval. Our framework also employs rigorous experimental design and validation, enabling us to select the most suitable model architecture and parameters to achieve optimal performance across multiple tasks.

Although PertOmni performs well across multiple datasets and tasks, it still has some limitations. First, the performance of PertOmni depends on the current LLM’s ability to integrate knowledge. Therefore, we need to track the progress of advanced LLM research to ensure we always use state-of-the-art embeddings. Second, PertOmni cannot directly process other informative modalities, such as protein-based perturbation data, due to data access limitation. By leveraging these multimodal perturbation datasets, we may push the unified setting of our current design. In the future, we will focus on these areas to continue improving the boundaries of PertOmni with the help of enriched datasets.

Overall, our work shows that LLM-derived priors can be combined with sequencing readouts to learn transferable perturbation representations. Future work could extend PertOmni to additional modalities and update LLM embeddings to improve generalization with larger and more diverse data.

## 4. Methods

### Problem definition

Our objective is to learn aligned representations of perturbation effects from single-cell transcriptomic perturbation data and LLM embeddings. Consider a perturbation dataset 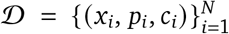, where *x*_*i*_ ∈ ℝ^*G*^ is the pseudobulk expression vector for sample *i, p*_*i*_ denotes the perturbation condition (gene knockdown or small molecule), and *c*_*i*_ denotes the corresponding cell line. We learn a transcriptome encoder *P*^*X*^ and text encoders *P*^*Z*^ that map inputs to a shared embedding space:

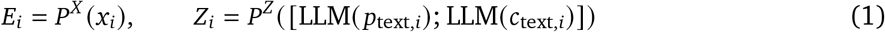

where 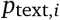 and 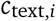 are textual descriptions of the perturbation and cell line for sample *i*, respectively, LLM(·) denotes a fixed text embedding model, [· ;·] denotes concatenation, and *E*_*i*_, *Z*_*i*_ ∈ ℝ^*d*^. We demonstrate that the learned transcriptome embeddings (*E*) and text embeddings (*Z*) support multiple downstream applications via linear probing or joint fine-tuning.

### Data preparation

For small-molecule perturbations, we use the Tahoe-100M dataset (8). For genetic perturbations, we use 5 cell lines across 3 datasets: K562 and RPE1 (40), HepG2 and Jurkat (9), and HCT116 (45). For both modalities, cells and genes are first filtered based on standard quality control criteria following the Scanpy tutorial (50), including the removal of cells that express fewer than 100 genes and the removal of genes that are detected in fewer than three cells. In each plate or batch, we perform pseudobulk aggregation by summing gene expression across cells sharing the same condition (perturbation and cell line). Then, normalization is performed following the Scanpy tutorial (50), where we count-normalize each pseudobulk sample to the median of total counts and apply a log1p transformation. For the Tahoe-100M dataset, text descriptions for perturbations are obtained directly from the dataset, where compound descriptions are extracted from MedChemExpress (51) using Selenium(52). For CRISPRi data, embeddings for target genes are taken from GenePT (43). For both datasets, cell line text descriptions are generated from GPT-4o using the prompt: “Please summarize the information of cell line: {cell line} in one paragraph. Use academic language and include pathway information.” All perturbation- and cell line-level text descriptions are then converted into 1536-dimensional embeddings using the text-embedding-3-small model. We split the training, validation, and test sets into 4/1/1 based on perturbations in the two datasets, ensuring that perturbations in the test set are not observed during training. For PertOmni-chem and PertOmni-cri training, we use all genes in Tahoe-100M dataset and CRISPRi dataset, respectively. For PertOmni training, we use the intersection of genes between the two datasets.

For the image-text retrieval task, we use the RxRx3-core dataset (48), a Cell Painting dataset curated from the RxRx3 dataset (18). The RxRx3-core dataset contains 222, 601 six-channel fluorescent microscopy images of HUVEC cells, covering 736 genetic perturbations and 1, 674 compound perturbations. We follow the image preprocessing steps from CellCLIP (47), where we normalize raw Cell Painting image values into 0 − 255. Then we treat each Cell Painting channel as an independent grayscale image and extract embeddings separately using pretrained DINOv2-giant model (49). For retrieval evaluation on unseen perturbations, we split the perturbations into 70 / 10 / 20 for training, validation and testing. For the text embedding, we first extract compound descriptions from MedChemExpress (51), following Tahoe-100M (8). For compounds that are not available on MedChemExpress, we generate compound text descriptions from GPT-4o using the prompt: “Please summarize the information of compound perturbation: {compound} in one paragraph. Use academic language and include pathway information.” Text descriptions for CRISPR perturbations and cell line are generated in the same manner. Then all text descriptions are converted into 1536-dimensional embeddings using the text-embedding-3-small model. The perturbation embeddings and cell line embeddings are concatenated to form the text embeddings for training PertOmni.

### Encoder architecture

All encoders 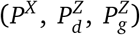 are lightweight projection heads with two fully-connected layers and a residual connection. Each maps an input of dimension *d* to a 128-dimensional latent vector: a linear layer to 128 dimensions, GELU, a second 128 × 128 linear layer with dropout, residual addition to the first projection, and layer normalization.

### Masked contrastive learning

Our model follows Contrastive Language-Image Pre-Training (CLIP) and its sigmoid-based variant SigLIP (54), aligning paired samples across modalities by increasing similarity of matched pairs and decreasing similarity of unmatched pairs. We also modify the masking process across different cell lines to learn better cellular representations. In our setting, each sample consists of a gene expression profile paired with LLM-derived embeddings of the perturbation and cell line.

We encode gene expression with a shared transcriptomic encoder *P*^*X*^ . We encode text with modality-specific text encoders: 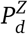 for small-molecule data and 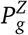 for genetic perturbation data. Each text encoder takes as input the concatenation of perturbation and cell-line text embeddings. For genetic perturbation data, we additionally concatenate LLM-derived embeddings of Gene Ontology biological pathways related to each target gene, where pathway descriptions are obtained from MSigDB (55, 56). We train *P*^*X*^ jointly with 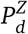 and 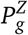 using a contrastive objective, and share the weights of the final fully-connected layer between 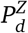 and 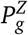 to encourage alignment across perturbation types. The architecture is visualized in Figure 1(a). We denote text embeddings for chemical perturbations as *T* and for genetic perturbations as *M*.

Let the small-molecule and genetic perturbation datasets contain *n* and *m* samples, respectively. Let *x*_*i*_ denote the input gene expression vector for sample *i*, and let *c*_text_, *p*_text_ be text descriptions of the cell line and perturbation. For the small-molecule perturbation dataset, we encode embeddings from the encoders as *E*_*i*_ = *P*^*X*^ (*x*_*i*_) and 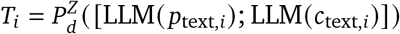 for the *i*_*th*_ sample. For the genetic perturbation dataset, encoded embeddings are 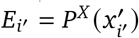 and 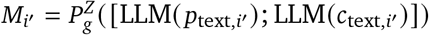 for the *i*^′^th sample. Our loss function is written as:

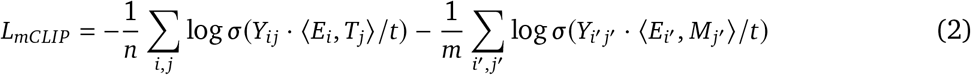

where *t* is temperature, *Y*_*i j*_, *Y*_*i*_′ _*j*_′ specify how each cross-modal pair contributes to the contrastive objective (positive, negative, or masked), and ⟨·, ·⟩ denotes cosine similarity. *σ* is the sigmoid function.

In standard SigLIP, *Y*_*i j*_ = +1 if *E*_*i*_ and *T*_*j*_ correspond to the same sample (matched pair) and *Y*_*i j*_ = − 1 otherwise (unmatched pair). In our setting, we perform contrastive learning only within cell lines. Concretely, we mask all pairs across different cell lines so that they do not contribute to the loss:

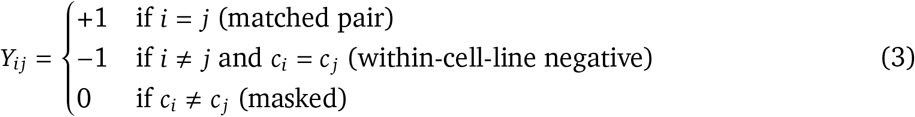

(and analogously for *Y*_*i*_′ _*j*_′ in the genetic perturbation dataset). Here, *c*_*i*_ denotes the discrete cell-line identity label used for masking. A visualization of *Y* is shown in Figure 1(b). Hyperparameters are described in Table 1.

**Table 1:**
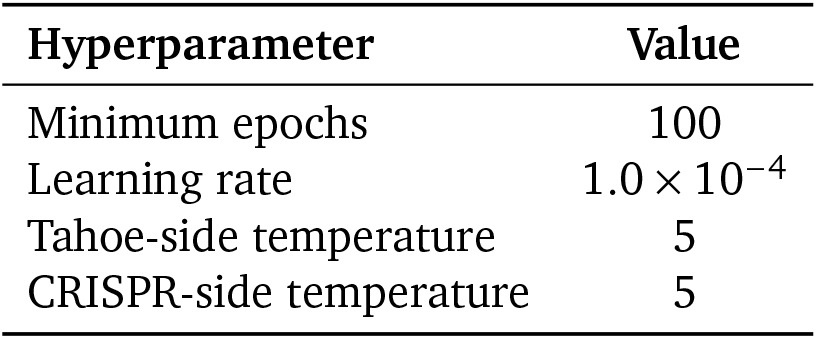
Training hyperparameters. Temperatures correspond to the scaling factor *t* in Eq. (2) for Tahoe (chemical) and CRISPR (genetic) datasets. Details of experiment settings are shown in Appendix A. Hyperparameter sensitivity is shown in Supplementary Figures.

| Hyperparameter | Value |
| --- | --- |
| Minimum epochs | 100 |
| Learning rate | $1.0 \times 10^{-4}$ |
| Tahoe-side temperature | 5 |
| CRISPR-side temperature | 5 |

**Table 2:**
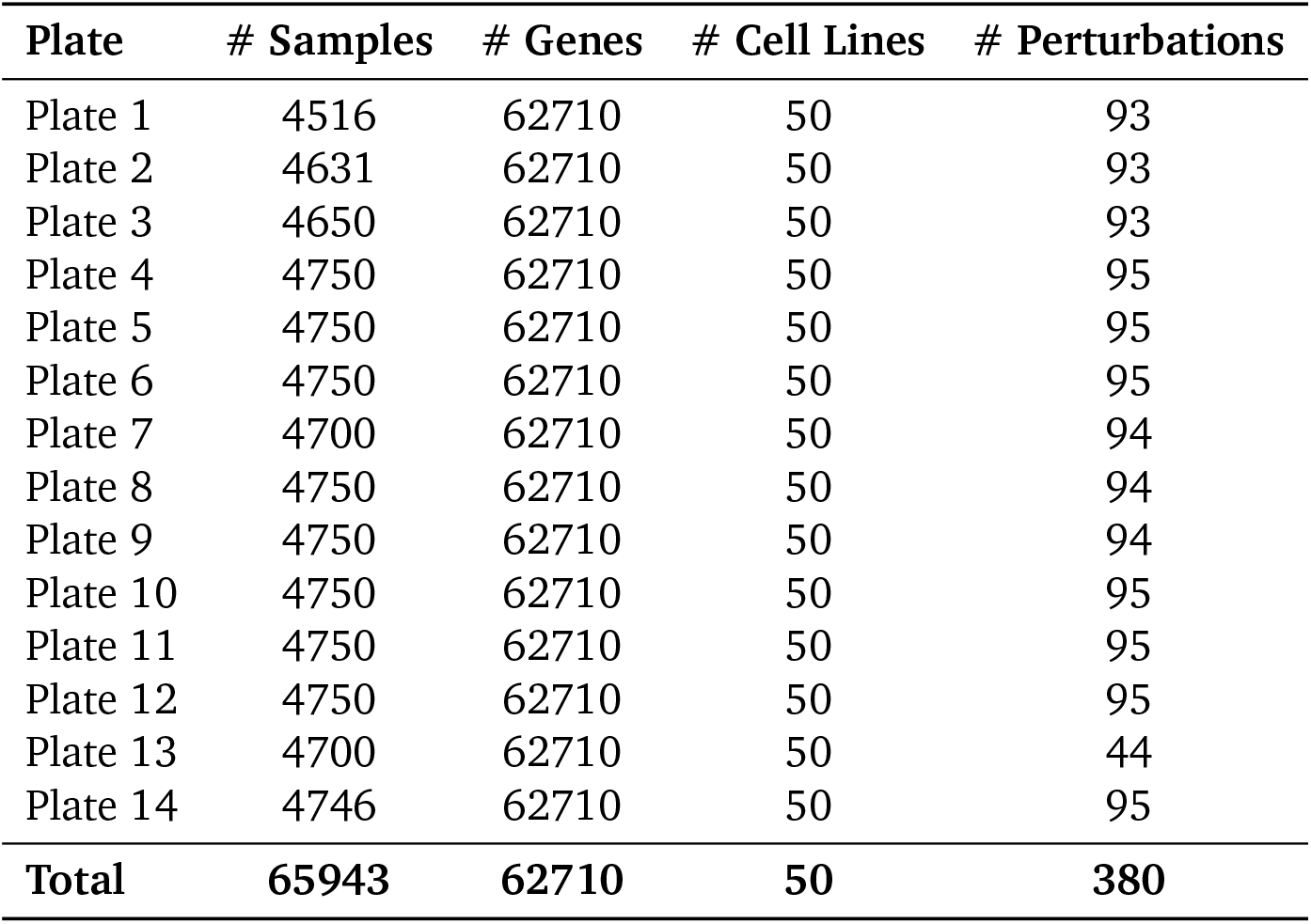
Summary statistics of the Tahoe-100M dataset across plates.

**Table 3:** Summary statistics of the CRISPR perturbation datasets across cell lines.

| Cell Line | # Samples | # Genes | # Cell Lines | # Perturbations |
| --- | --- | --- | --- | --- |
| K562 | 11818 | 6408 | 1 | 9783 |
| RPE1 | 2373 | 6408 | 1 | 2373 |
| HepG2 | 2373 | 6408 | 1 | 2373 |
| Jurkat | 2373 | 6408 | 1 | 2373 |
| HCT116 | 18115 | 6408 | 1 | 18115 |
| **Total** | **37052** | **6408** | **5** | **18385** |

**Table 4:**
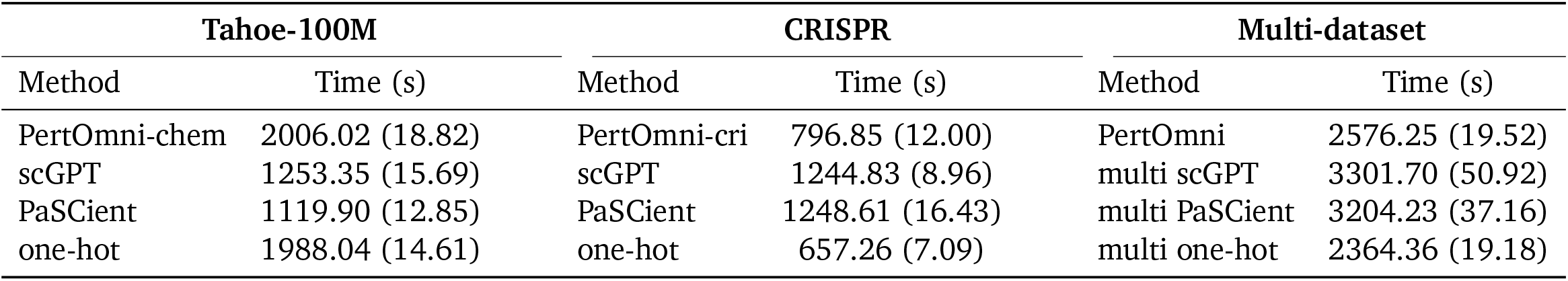
Runtime (seconds; mean (std)) on Tahoe-100M, CRISPR, and multi-dataset settings. We use 8 CPUs and 1 NVIDIA RTX 5000 GPU for this experiment.

Cross–cell-line negatives in standard contrastive learning can encourage the model to separate samples primarily by cell identity rather than perturbation identity, because baseline expression programs differ strongly across contexts. By setting *Y*_*i j*_ = 0 when *c*_*i*_ ≠ *c* _*j*_ (Eq. (3)), we prevent the objective from using cell line as an easy shortcut and instead force discrimination among perturbations within the same context. This design makes the learned space well-suited for context-conditional tasks (e.g., retrieval and kNN prediction within a given cell line), at the cost of not explicitly enforcing that a perturbation’s embedding be invariant across cell lines. In joint training, cross-context generalization can still emerge indirectly through shared encoder parameters and shared perturbation text priors, but we do not assume perturbation effects are globally comparable across cell types.

### Perturbation retrieval

Using the learned transcriptome and text embeddings from PertOmni, we perform intra-cell-line retrieval between different modalities based on cosine similarity. For this task, we evaluate retrieval performance using three retrieval metrics shown below:

Let *Q* be a set of queries. For each query *q* ∈ *Q*, let *g*_*q*_ be the ground-truth target and 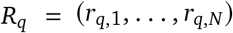 be the ranked list of retrieved items. Define *rank*_*q*_ = min{*i* ∈ {1, …, *N*} | *r*_*q,i*_ = *g*_*q*_}.

1. Accuracy score for top-K retrieval measures whether the ground truth appears within the retrieved candidates. It is defined as

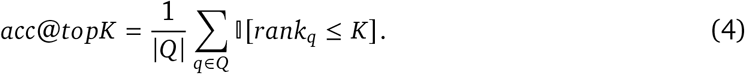
2. Normalized discounted cumulative gain (nDCG) score for top-K retrieval measures overall ranking quality. It is defined as

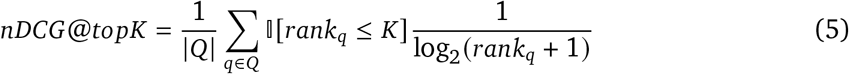
3. Mean reciprocal rank (MRR) score captures how early the ground-truth target appears in the ranked list. It is defined as

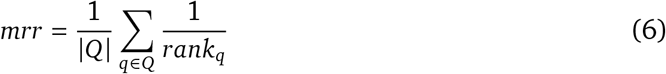

### Gene-drug interaction prediction

This task aims to identify target genes for chemical perturbations (Drug2Gene) and identify chemicals that target a specific gene (Gene2Drug). We fine-tune PertOmni using interaction annotations from DGIdb (57). Since DGIdb is incomplete, the absence of an annotation does not imply a true negative interaction. We therefore treat this as a positive– unlabeled learning problem and optimize a positive-only objective: annotated drug–gene pairs are encouraged to have high similarity in the learned embedding space, while unannotated pairs are ignored during the development process.

Let *A*_*i j*_ ∈ {0, 1} denote whether DGIdb annotates an interaction between drug *i* and gene *j*, and let *s*_*i j*_ = ⟨*T*_*i*_, *M*_*j*_⟩ / *t* be the scaled cosine similarity between the corresponding embeddings. To train the model, we optimize the following objective:

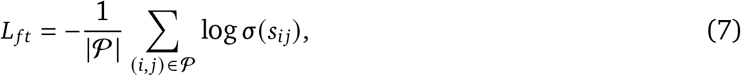

where *P* = {(*i, j*) : *A*_*i j*_ = 1}, *T*_*i*_ and *M* _*j*_ are outputs from the drug and gene text encoders of PertOmni, respectively.

For this task (PertOmni_*ft*_), we initialize from the pretrained PertOmni model and optimize only the dataset-specific text encoders 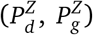 under Eq. (7), while keeping the shared transcriptome encoder (*P*^*X*^ ) fixed.

Inspired by (36), we also consider training a Siamese neural network (SNN) on transcriptomic, text, and all (transcriptomic+text) embedding as a strong baseline for comparison. The SNN applies shared weights to a pair of drug and gene embeddings, followed by element-wise multiplication of the resulting representations and a classifier that predicts the probability of a drug-gene pairing. Following (36), we apply a balanced training strategy to prevent the SNN from being biased towards well-known targets. Specifically, for the Drug2Gene task, for each gene in a positive pair, we sample a drug that is not annotated to target that gene to form an unannotated contrast pair; for the Gene2Drug task, the other way round.

For evaluation, we use AUROC and AUPRC as metrics, since this is a binary classification problem for each drug-gene pair. Here we treat unannotated pairs as negatives, noting this likely underestimates performance due to label incompleteness.

### Perturbation prediction

Inspired by LangPert (39), we perform retrieval between text embedding in the testing set and text or transcriptome embedding in the training set, followed by k-nearest neighbors (kNN) aggregation of retrieved training samples’ expression profiles to generate predictions. Additionally, we first use retrieval to narrow down candidate perturbations, then use the refined LangPert model to obtain the final prediction. Model performance is evaluated using Pearson correlation between predicted and observed gene expression, computed over all genes or the top 20 differentially expressed genes (DEGs).

### Transcriptomic embedding ablations

We perform an ablation study to evaluate alternative methods for embedding transcriptomic profiles. Either pseudobulk or embeddings are then used as input *x* to the transcriptome encoder *P*^*X*^ in PertOmni. The evaluation uses cross-modality retrieval as a metric. We compare PertOmni directly on pseudobulk with two foundation-model baselines: scGPT (27) and PaSCient (46). We generate cell-level embeddings using scGPT, then aggregate them to the sample level, whereas PaSCient directly generates sample-level embeddings. In addition, we test random matching and one-hot embeddings in place of the LLM-derived perturbation and cell line embeddings. Training cost comparisons are presented in Supplementary Table 4.

### Image-text retrieval

We follow CellCLIP (47) and extract a frozen 1536-dimensional DINOv2 embedding for each channel, resulting in five channel-wise embeddings per image. We considered three channel aggregation strategies. Concat_PertOmni concatenates the five channel-wise embeddings into a 7680-dimensional vector. ProjMean_PertOmni applies a shared MLP projection head to each channel embedding and then mean-pools the projected embeddings across channels. Mean_PertOmni directly averages the five channel-wise embeddings before projection. Image embeddings are averaged to the perturbation level before contrastive training and retrieval evaluation. Using the image and text embeddings, We train the same model architecture and contrastive loss as previous transcriptome-text retrieval task. We compare PertOmni with the pretrained CellCLIP checkpoint and a CellCLIP model fine-tuned on the RxRx3-core dataset.

## 5. Ethics Statement and Acknowledgment

The authors used LLMs to generate fixed text embeddings for model training and editing of the manuscript, such as rephrasing for clarity.

## A. Experimental settings

For model training, we utilize 128 as batch size, 1e-4 as learning rate, Adam as optimizer, 0.0 as dropout rate, 100 as default epochs, 5.0 as temperature, and top-1 retrieval accuracy as the criteria for early stopping. The choices of hyper parameters satisfy for both cross-modality retrieval and drug-gene interaction prediction tasks.

## B. Supplementary Tables

## C. Supplementary Figures

**Supplementary Figure 1:**
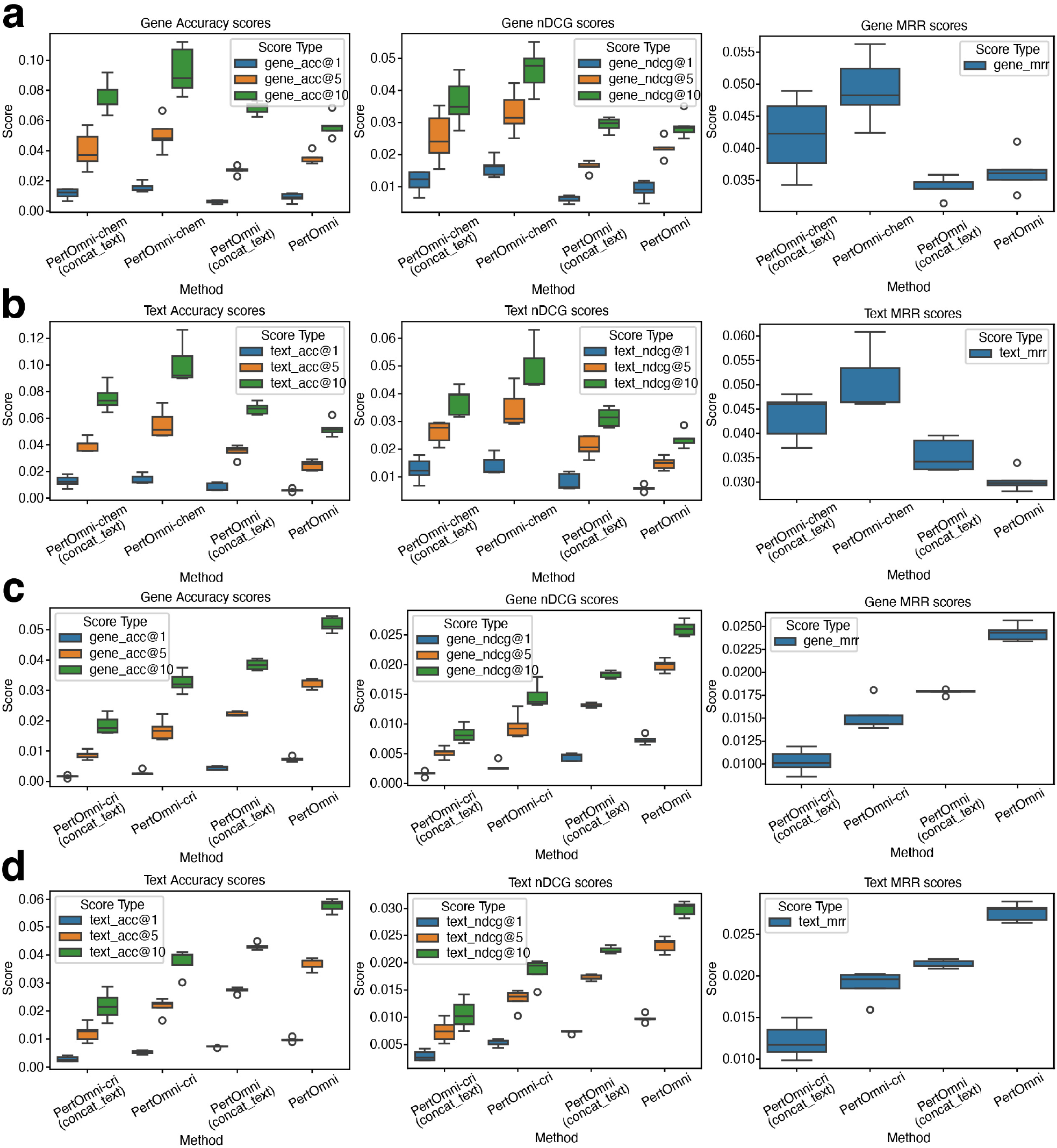
Results of cross-modality retrieval in ablation studies. In this ablation study, we first concatenate text descriptions of perturbation and cell line, and then generate LLM embeddings based on the concatenated descriptions. In contrast, PertOmni first generates LLM embeddings for perturbation and cell line, respectively, and then concatenates LLM embeddings as input to text encoders. (a) Gene2Text retrieval metrics comparison based on the Tahoe-100M dataset. (b) Text2Gene retrieval metrics comparison based on the merged CRISPR dataset.

**Supplementary Figure 2:**
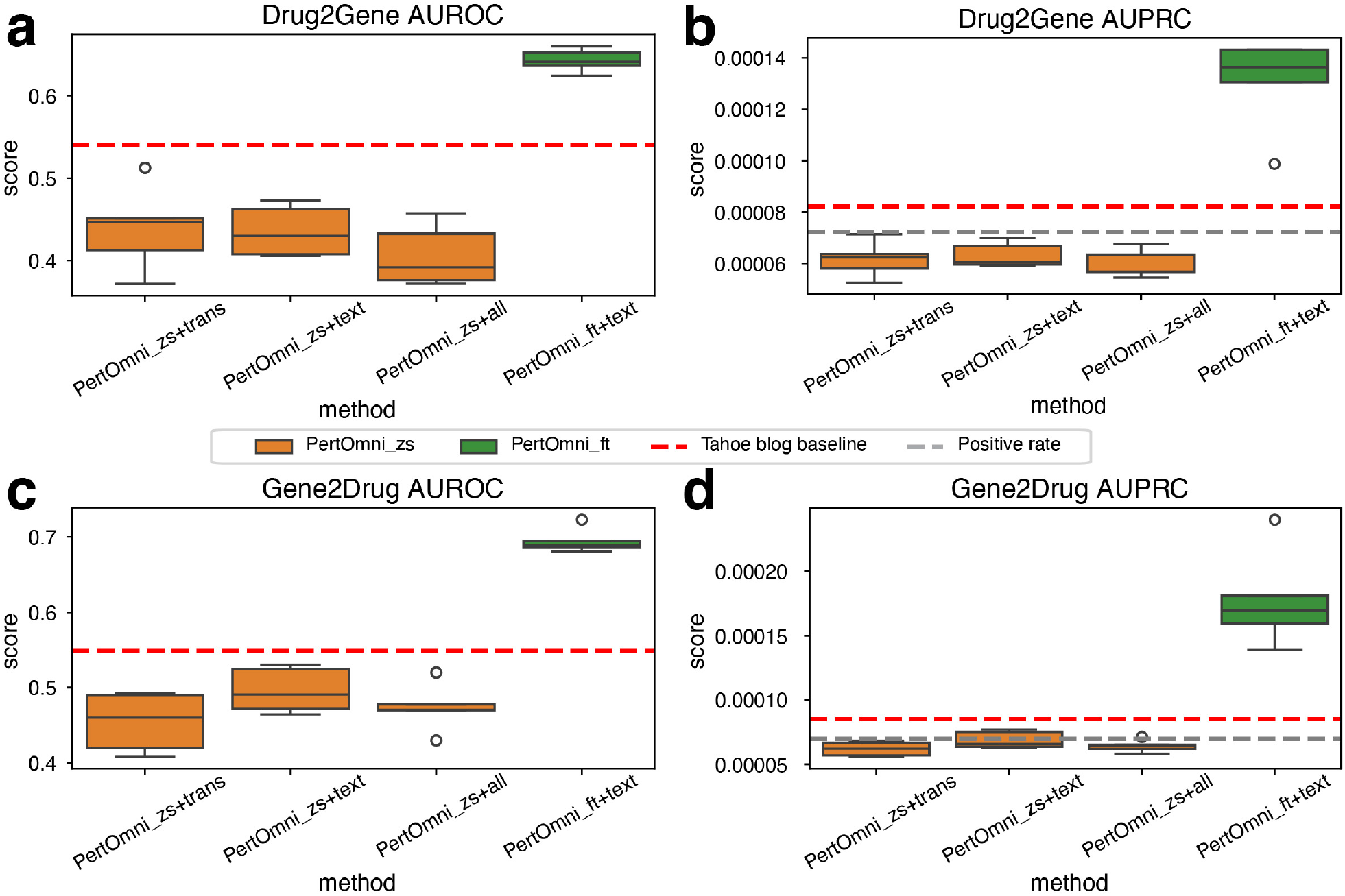
Comparison of drug-gene interaction performance before and after finetuning. PertOmni_zs represents the zero-shot performance of PertOmni without training based on paired gene-drug interactions, and PertOmni_ft represents fine-tuned performance. We find that the finetuning (ft) mode performs better. (a) AUROC comparison for the Drug2Gene task. (b) AUPRC comparison for the Drug2Gene task. (c) AUROC comparison for the Gene2Drug task. (d) AUPRC comparison for the Gene2Drug task.

**Supplementary Figure 3:**
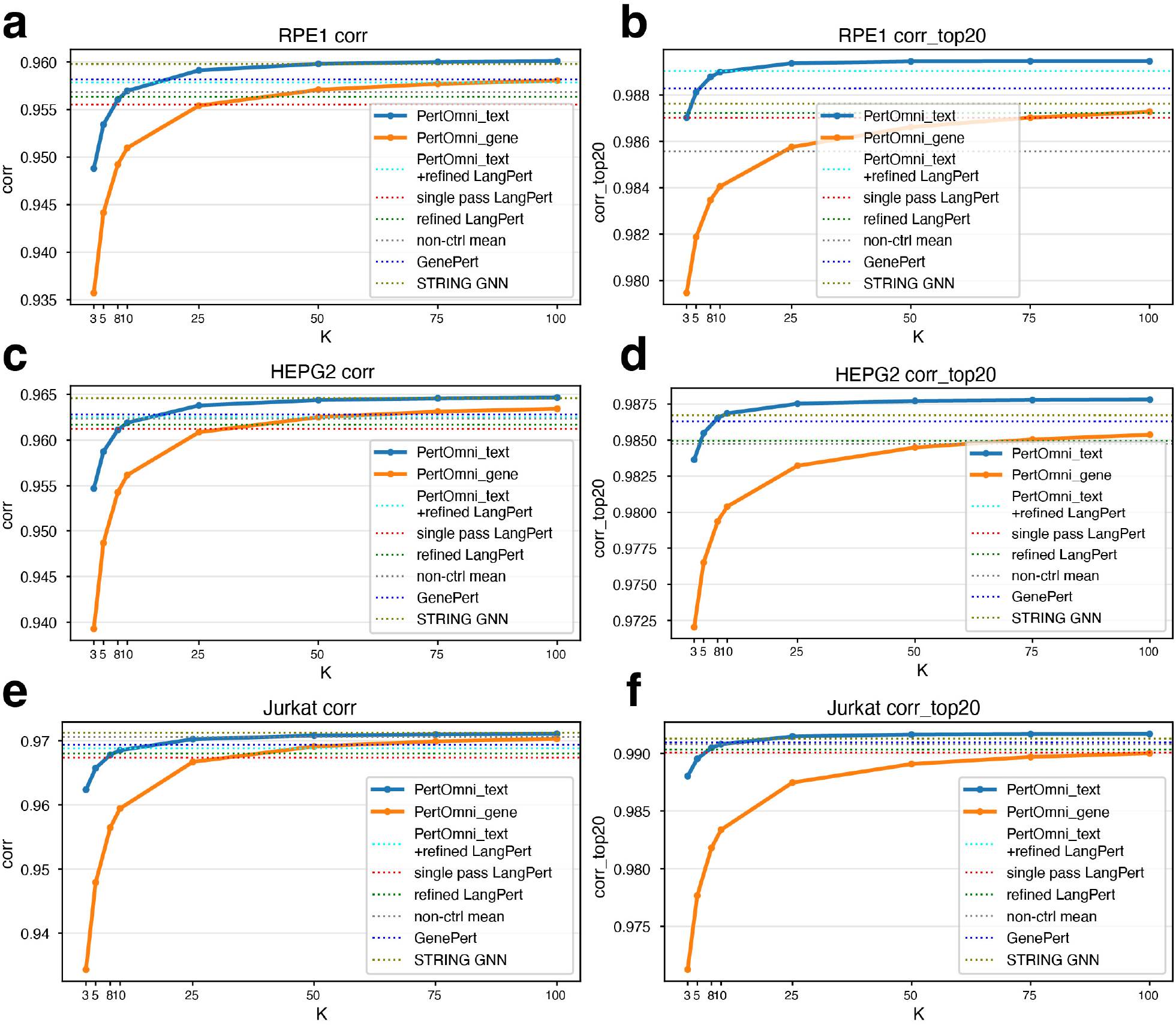
Linechart visualization of how perturbation prediction performance changes with *k*. a) All genes, RPE1. (b) Top-20 DEGs, RPE1. (c) All genes, HepG2. (d) Top-20 DEGs, HepG2. (e) All genes, Jurkat. (f) Top-20 DEGs, Jurkat.

**Supplementary Figure 4:**
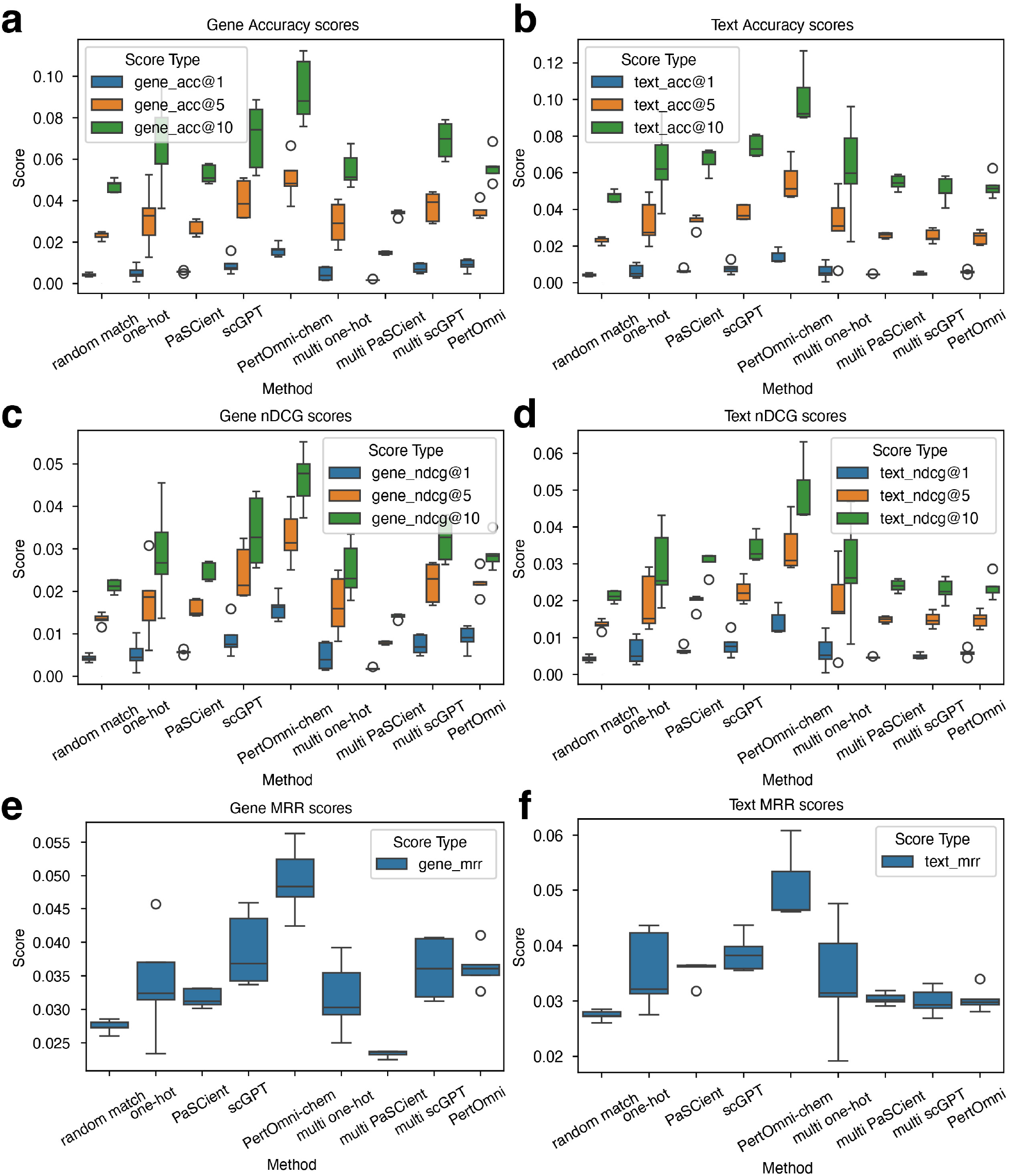
Boxplot visualization of cross-modality retrieval in Tahoe-100m dataset. (a,b) Accuracy retrieval scores comparison. (c,d) nDCG scores comparison. (e,f) MRR scores comparison.

**Supplementary Figure 5:**
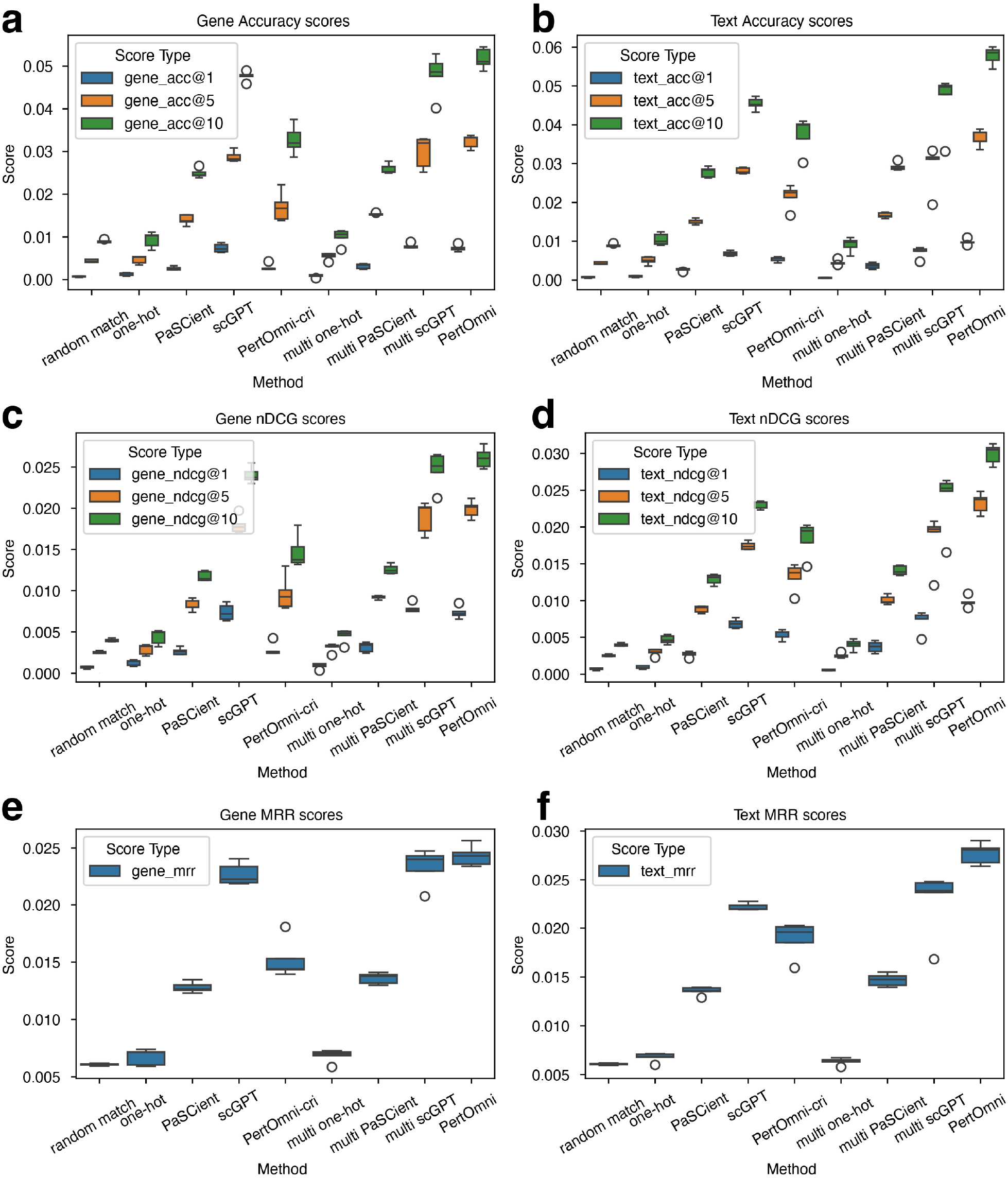
Boxplot visualization of cross-modality retrieval in the CRISPR dataset. (a,b) Accuracy retrieval scores comparison. (c,d) nDCG scores comparison. (e,f) MRR scores comparison.

**Supplementary Figure 6:**
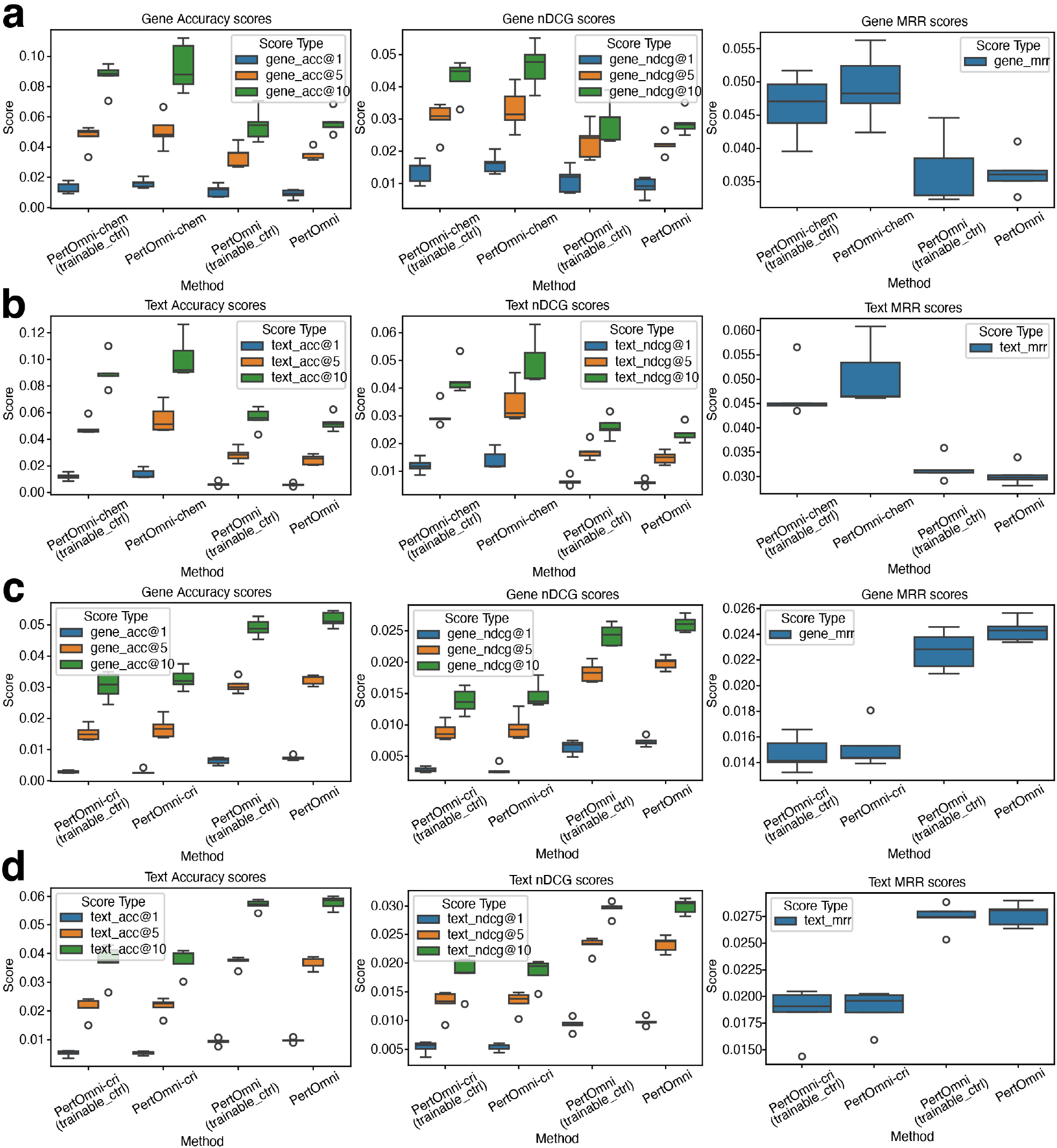
Results of cross-modality retrieval in ablation studies. In this ablation study, we use trainable embedding as input to text encoders for control samples (“DMSO_TF” in Tahoe-100M dataset and “non-targeting” in CRISPR dataset), instead of using 0s as in PertOmni. (a) Gene2Text retrieval metrics comparison based on the Tahoe-100M dataset. (b) Text2Gene retrieval metrics comparison based on the merged CRISPR dataset.

**Supplementary Figure 7:**
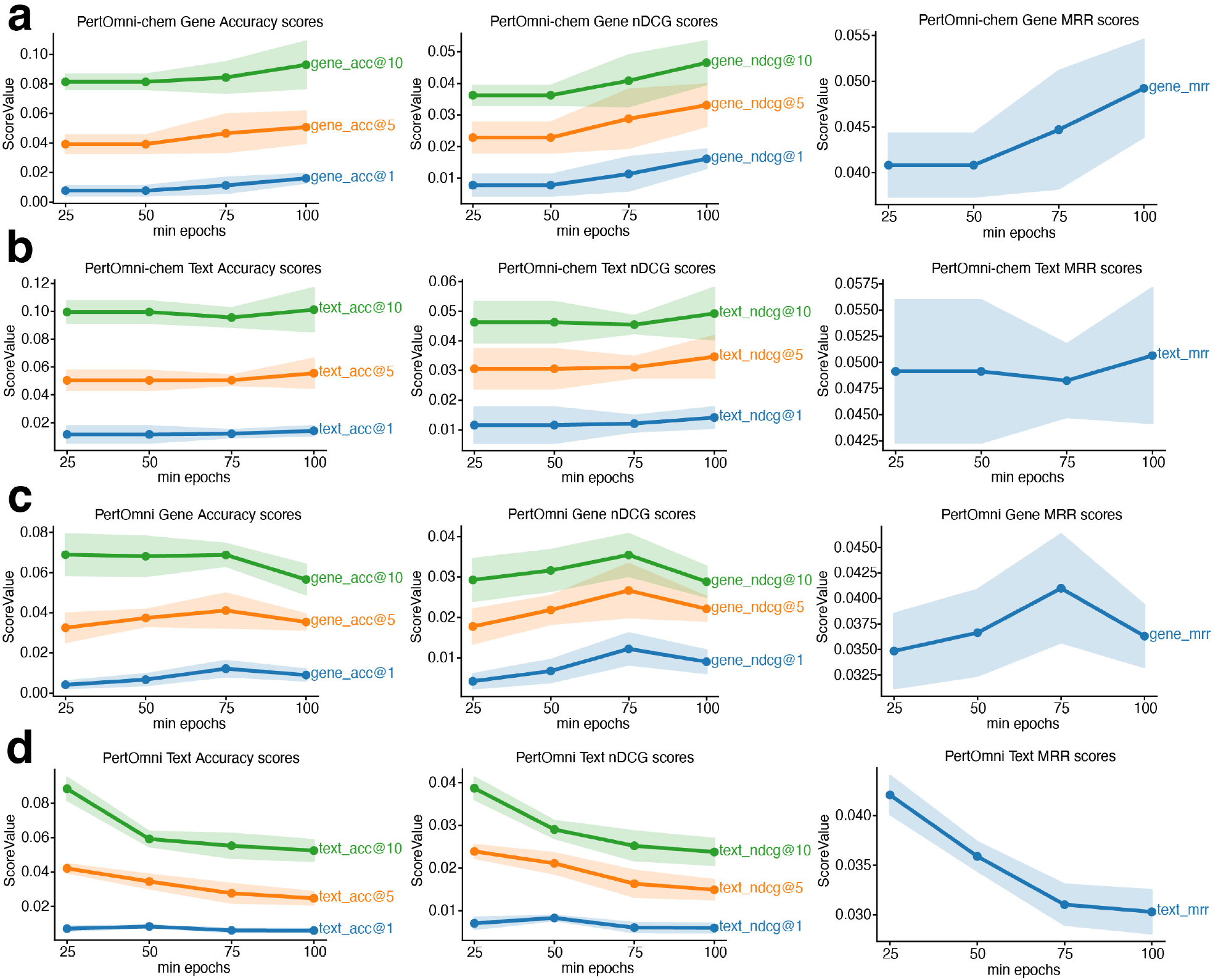
Results of cross-modality retrieval under different minimum epochs settings in Tahoe-100m dataset. (a) Gene2Text retrieval metrics comparison for PertOmni-chem. (b) Text2Gene retrieval metrics comparison for PertOmni-chem. (c) Gene2Text retrieval metrics comparison for PertOmni. (d) Text2Gene retrieval metrics comparison for PertOmni.

**Supplementary Figure 8:**
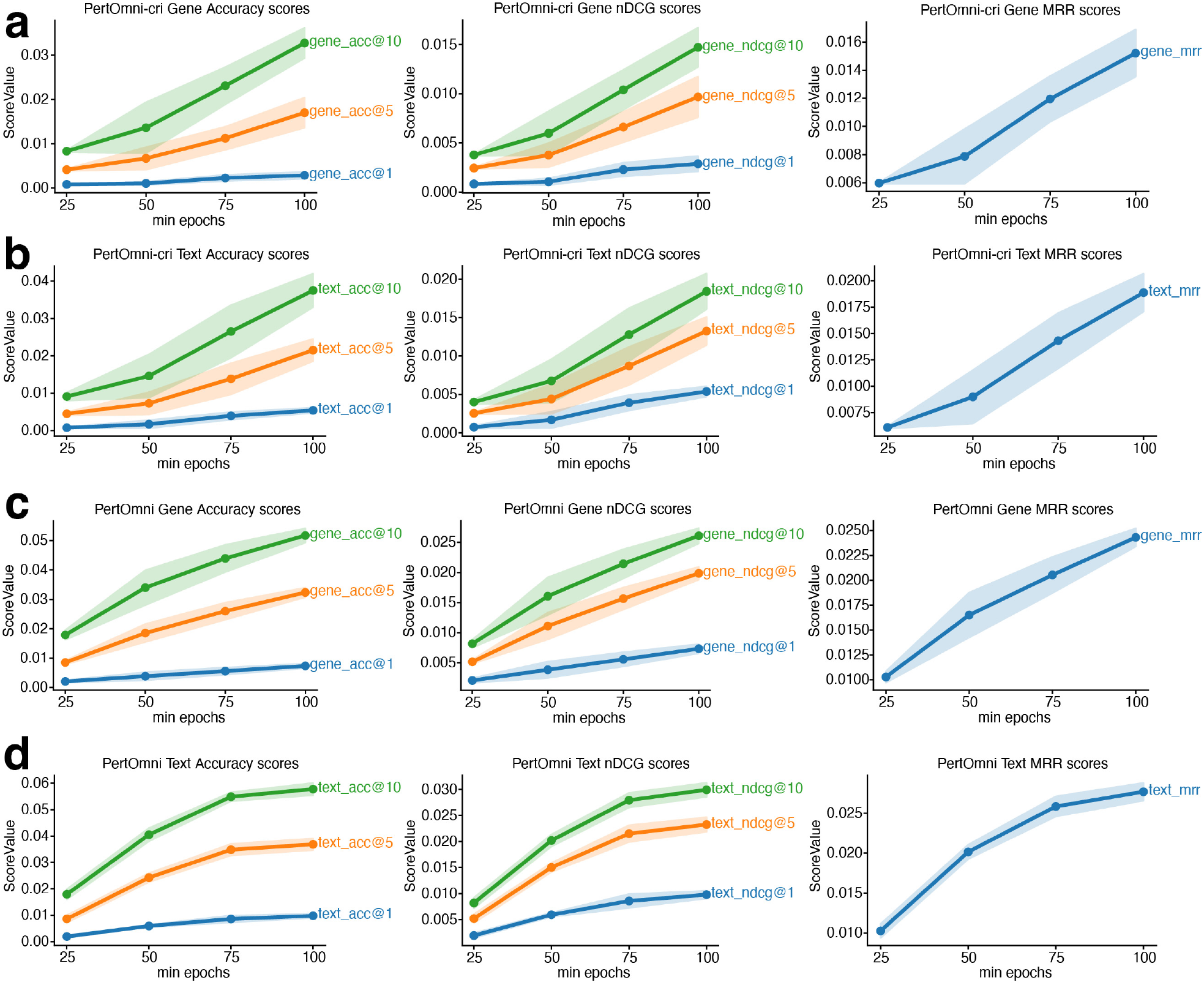
Results of cross-modality retrieval under different minimum epochs settings in the CRISPR dataset. (a) Gene2Text retrieval metrics comparison for PertOmni-cri. (b) Text2Gene retrieval metrics comparison for PertOmni-cri. (c) Gene2Text retrieval metrics comparison for PertOmni. (d) Text2Gene retrieval metrics comparison for PertOmni.

**Supplementary Figure 9:**
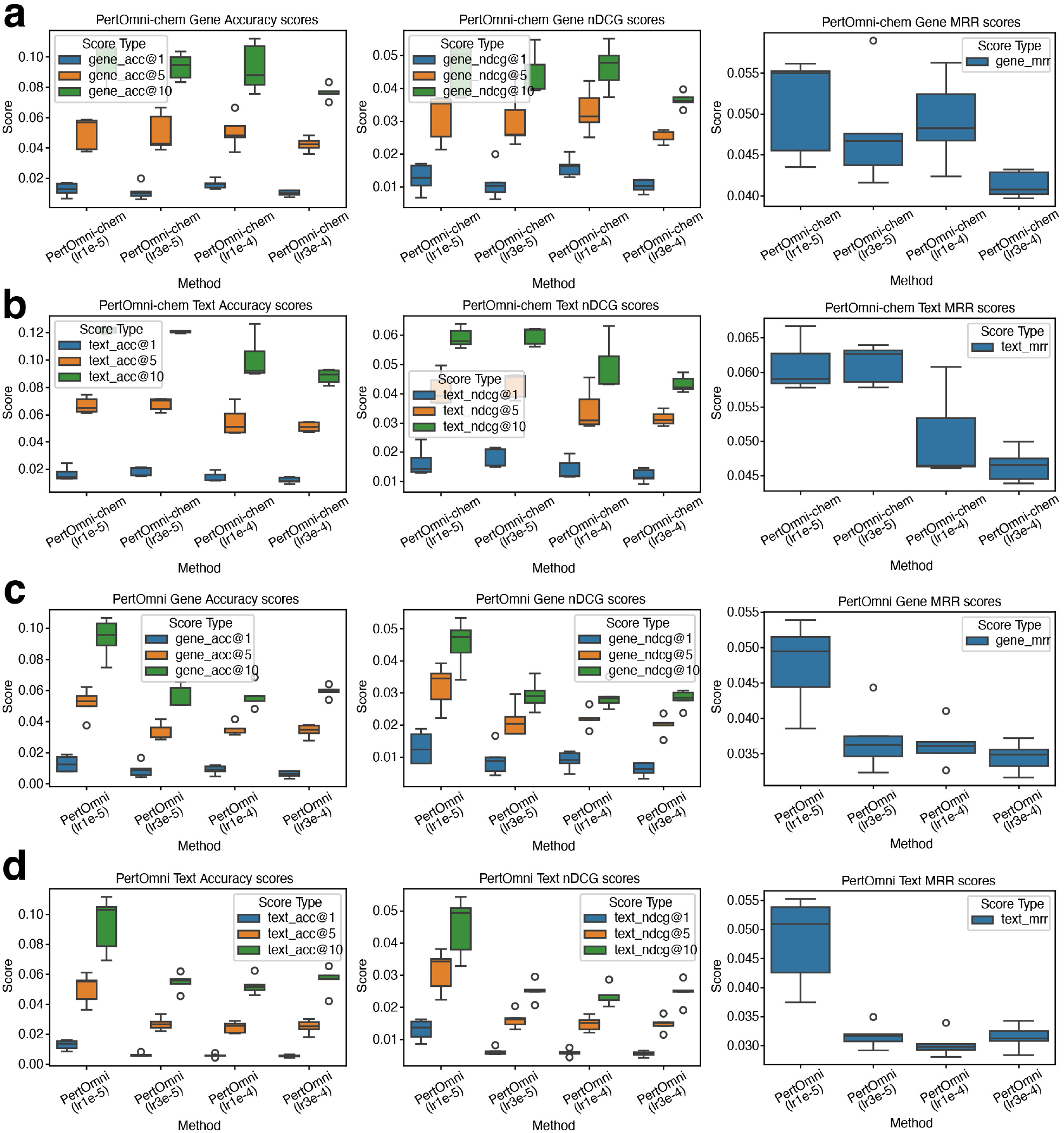
Results of cross-modality retrieval under different learning rate settings in Tahoe-100m dataset. (a) Gene2Text retrieval metrics comparison for PertOmni-chem. (b) Text2Gene retrieval metrics comparison for PertOmni-chem. (c) Gene2Text retrieval metrics comparison for PertOmni. (d) Text2Gene retrieval metrics comparison for PertOmni.

**Supplementary Figure 10:**
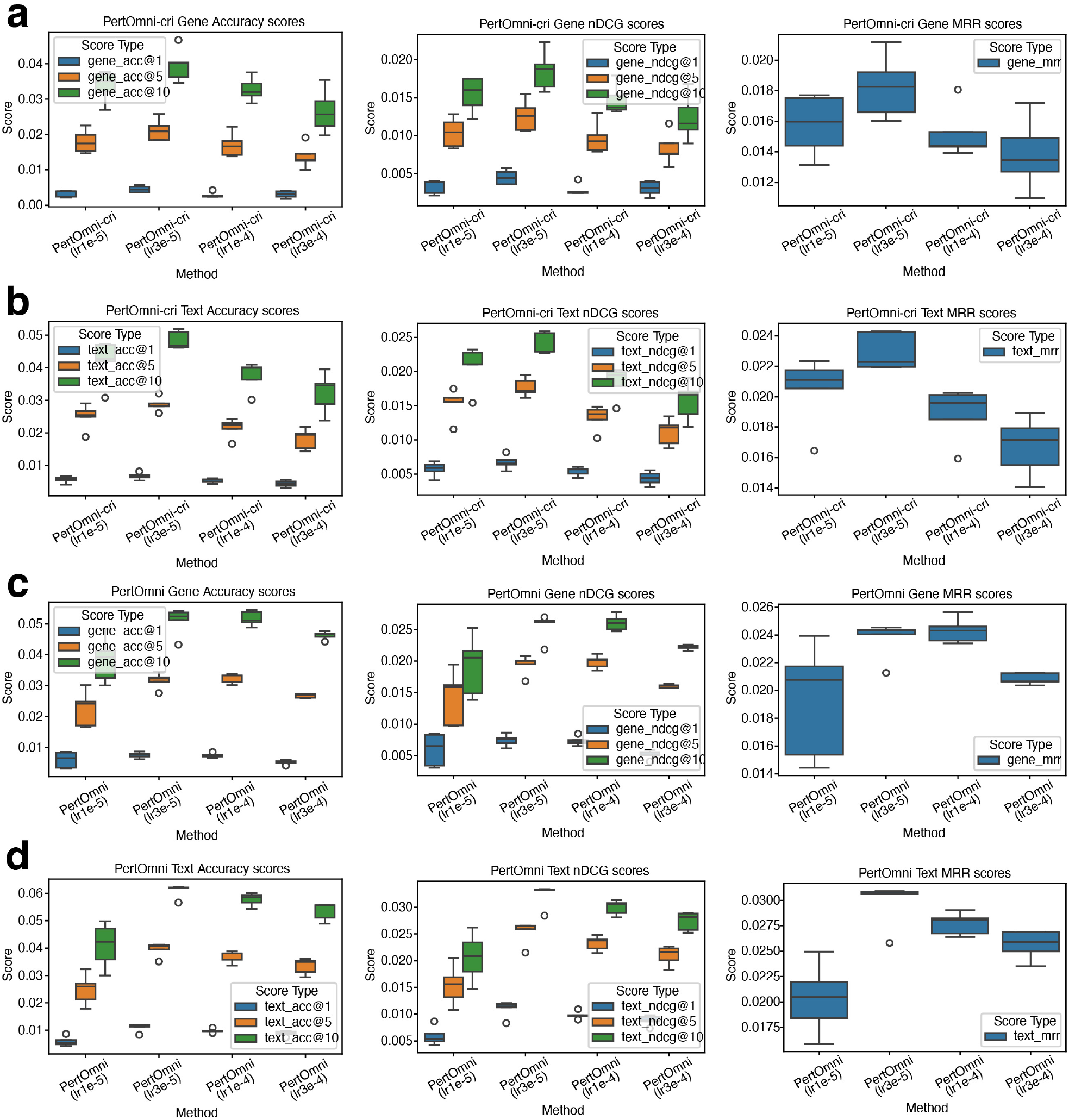
Results of cross-modality retrieval under different learning rate settings in the CRISPR dataset. (a) Gene2Text retrieval metrics comparison for PertOmni-cri. (b) Text2Gene retrieval metrics comparison for PertOmni-cri. (c) Gene2Text retrieval metrics comparison for PertOmni. (d) Text2Gene retrieval metrics comparison for PertOmni.

**Supplementary Figure 11:**
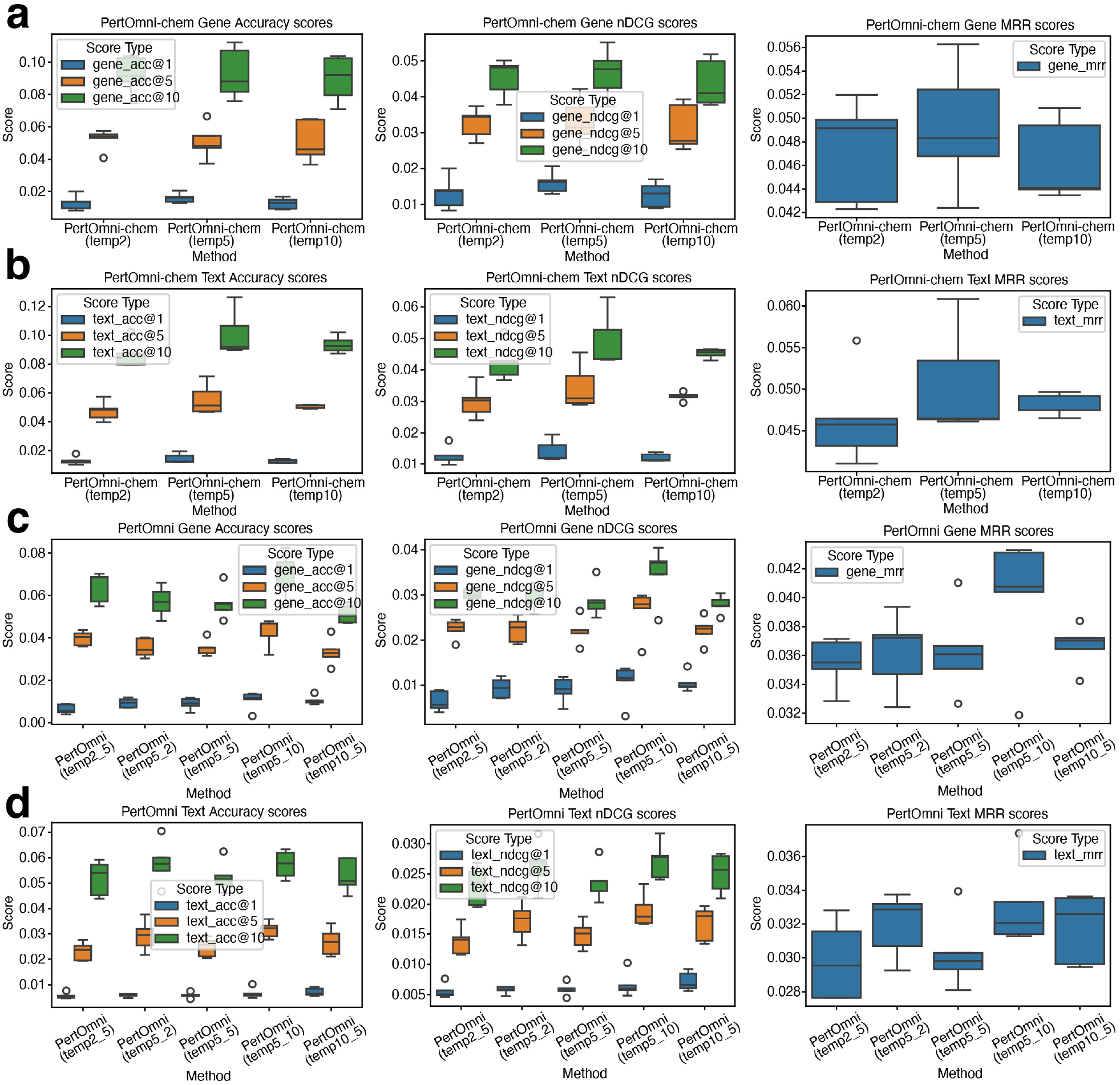
Results of cross-modality retrieval under different temperature settings in the Tahoe-100m dataset. (a) Gene2Text retrieval metrics comparison for PertOmni-chem. (b) Text2Gene retrieval metrics comparison for PertOmni-chem. (c) Gene2Text retrieval metrics comparison for PertOmni. (d) Text2Gene retrieval metrics comparison for PertOmni.

**Supplementary Figure 12:**
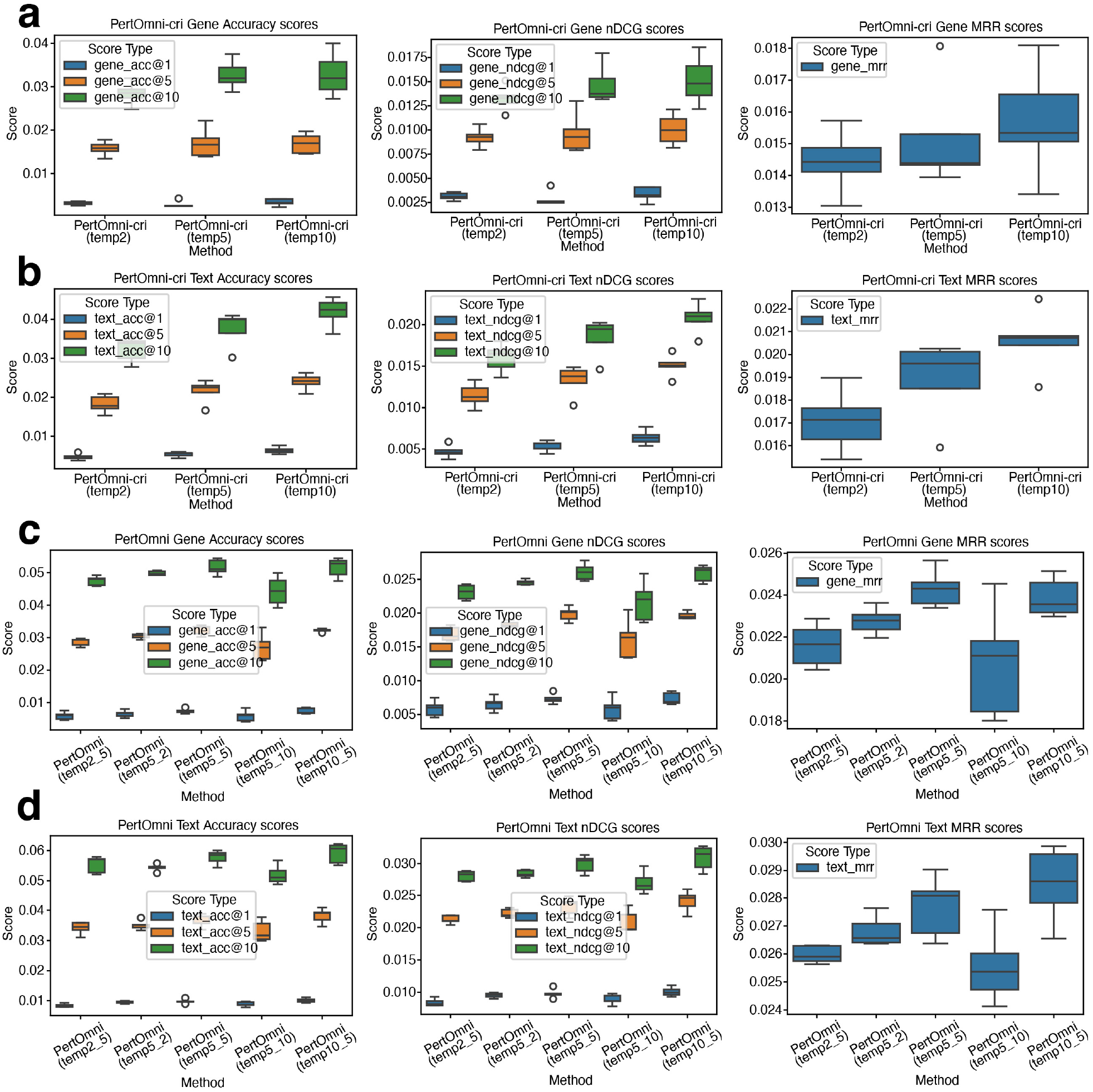
Results of cross-modality retrieval under different temperature settings in the CRISPR dataset. (a) Gene2Text retrieval metrics comparison for PertOmni-cri. (b) Text2Gene retrieval metrics comparison for PertOmni-cri. (c) Gene2Text retrieval metrics comparison for PertOmni. (d) Text2Gene retrieval metrics comparison for PertOmni.

**Supplementary Figure 13:**
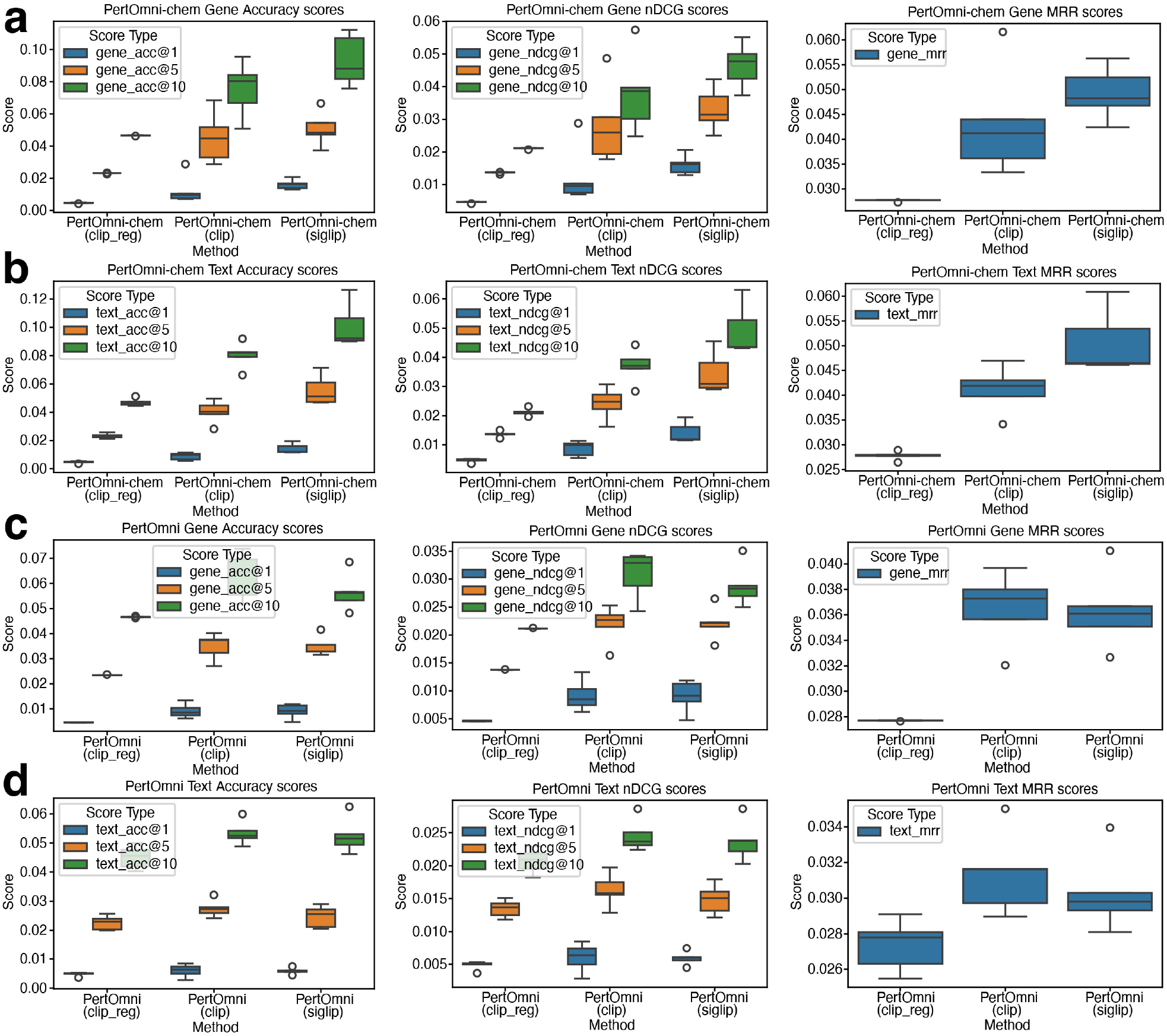
Results of cross-modality retrieval under different loss functions in the Tahoe-100m dataset. (a) Gene2Text retrieval metrics comparison for PertOmni-chem. (b) Text2Gene retrieval metrics comparison for PertOmni-chem. (c) Gene2Text retrieval metrics comparison for PertOmni. (d) Text2Gene retrieval metrics comparison for PertOmni.

**Supplementary Figure 14:**
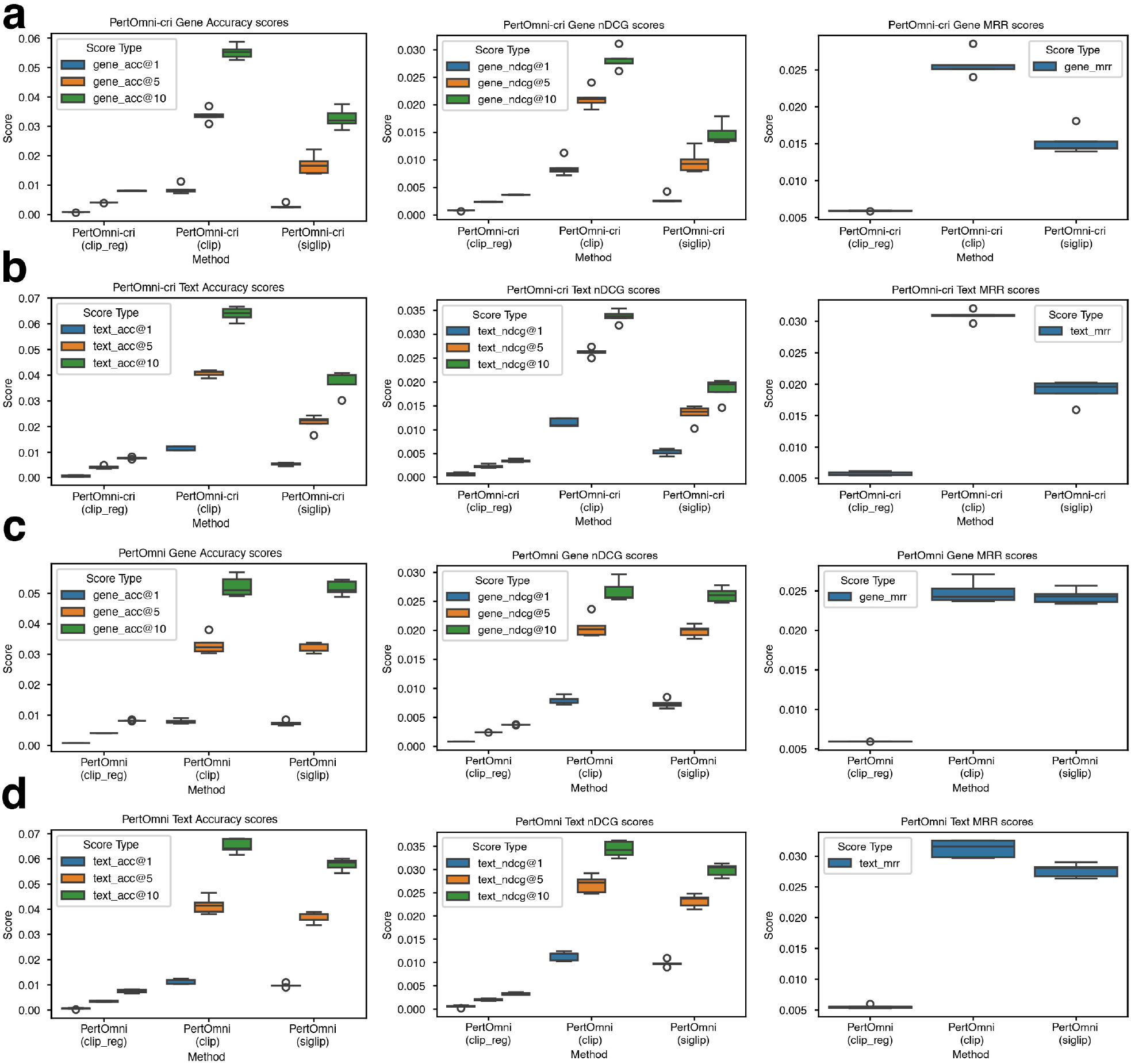
Results of cross-modality retrieval under different loss functions in the CRISPR dataset. (a) Gene2Text retrieval metrics comparison for PertOmni-cri. (b) Text2Gene retrieval metrics comparison for PertOmni-cri. (c) Gene2Text retrieval metrics comparison for PertOmni. (d) Text2Gene retrieval metrics comparison for PertOmni.

**Supplementary Figure 15:**
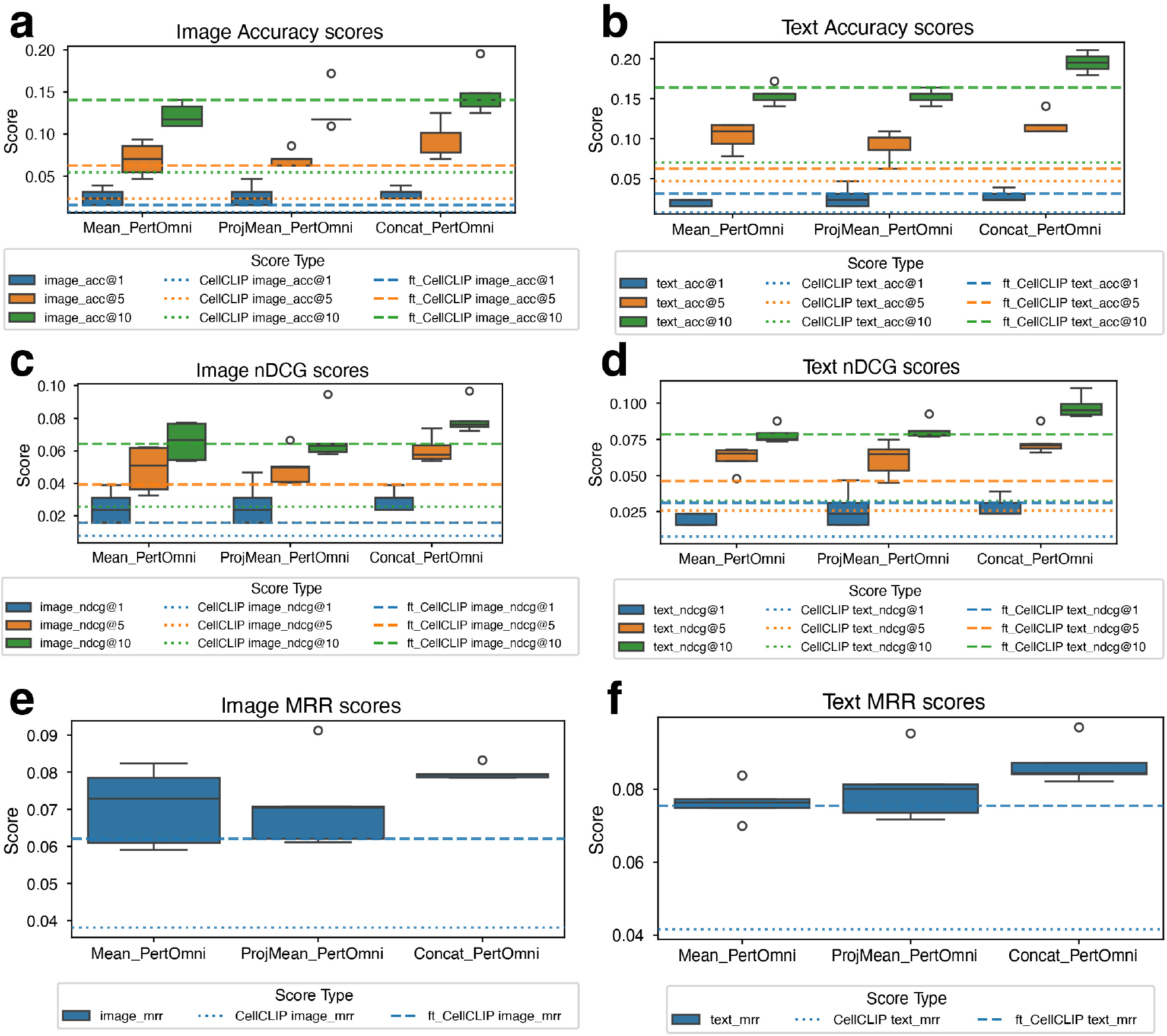
Boxplot visualization of cross-modality retrieval in compound perturbations of RxRx3-core dataset. (a,b) Accuracy retrieval scores comparison. (c,d) nDCG scores comparison. (e,f) MRR scores comparison.

**Supplementary Figure 16:**
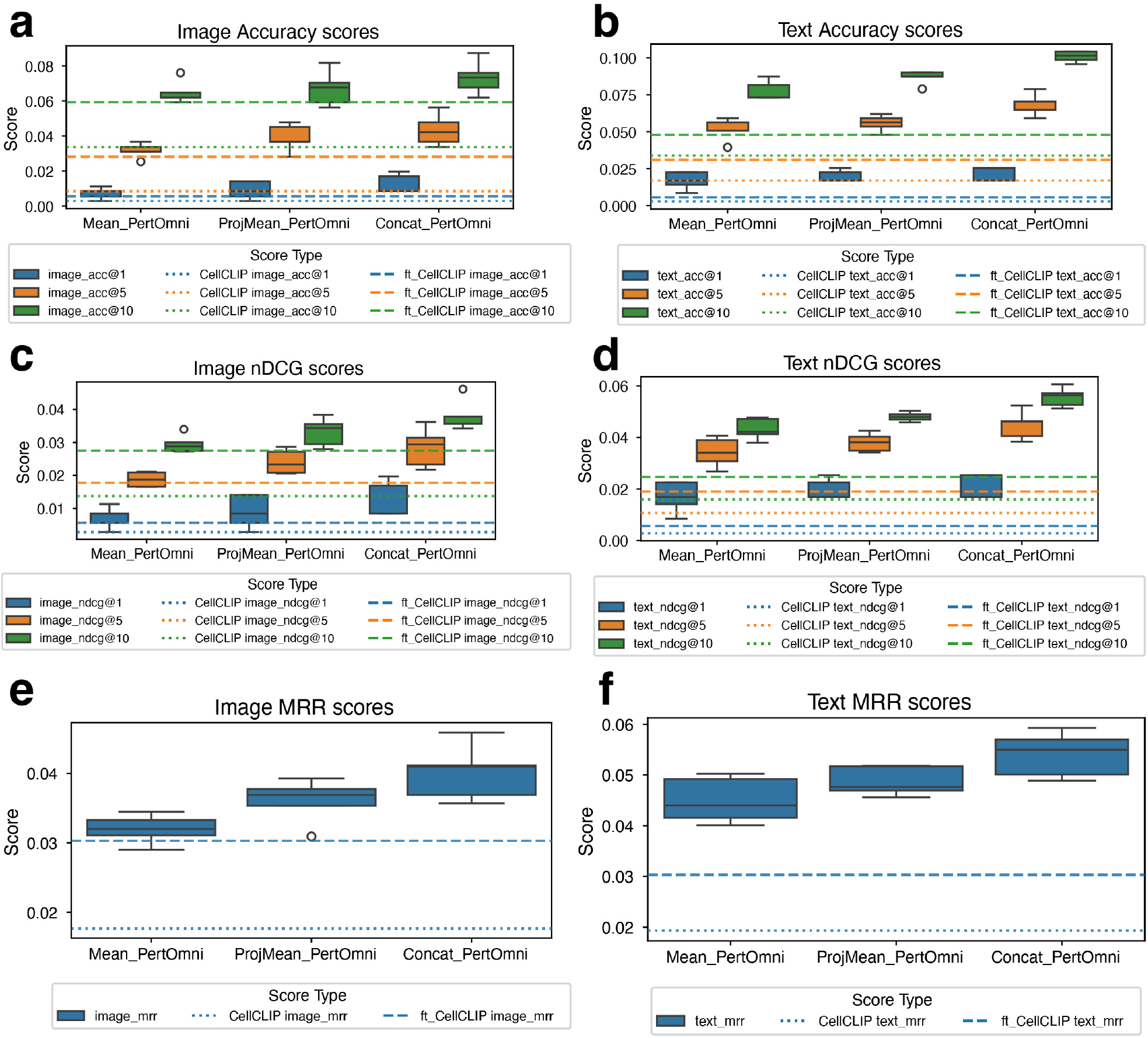
Boxplot visualization of cross-modality retrieval in crispr perturbations of RxRx3-core dataset. (a,b) Accuracy retrieval scores comparison. (c,d) nDCG scores comparison. (e,f) MRR scores comparison.

## Footnotes

1 Codebase: https://anonymous.4open.science/r/PertOmni-D728/

## References

[1] Paul Datlinger, André F Rendeiro, Christian Schmidl, Thomas Krausgruber, Peter Traxler, Johanna Klughammer, Linda C Schuster, Amelie Kuchler, Donat Alpar, and Christoph Bock. Pooled crispr screening with single-cell transcriptome readout. Nature Methods, 14(3):297–301, 2017.

[2] Atray Dixit, Oren Parnas, Biyu Li, Jenny Chen, Charles P Fulco, Livnat Jerby-Arnon, Nemanja D Marjanovic, Danielle Dionne, Tyler Burks, Raktima Raychowdhury, et al. Perturb-seq: dissecting molecular circuits with scalable single-cell rna profiling of pooled genetic screens. Cell, 167(7):1853–1866, 2016.

[3] Diego Adhemar Jaitin, Assaf Weiner, Ido Yofe, David Lara-Astiaso, Hadas Keren-Shaul, Eyal David, Tomer Meir Salame, Amos Tanay, Alexander van Oudenaarden, and Ido Amit. Dissecting immune circuits by linking crispr-pooled screens with single-cell rna-seq. Cell, 167(7):1883–1896, 2016.

[4] Luke A Gilbert, Max A Horlbeck, Britt Adamson, Jacqueline E Villalta, Yuwen Chen, Evan H Whitehead, Carla Guimaraes, Barbara Panning, Hidde L Ploegh, Michael C Bassik, et al. Genome-scale crispr-mediated control of gene repression and activation. Cell, 159(3):647–661, 2014.

[5] Britt Adamson, Thomas M Norman, Marco Jost, Min Y Cho, James K Nuñez, Yuwen Chen, Jacqueline E Villalta, Luke A Gilbert, Max A Horlbeck, Marco Y Hein, et al. A multiplexed single-cell crispr screening platform enables systematic dissection of the unfolded protein response. Cell, 167(7):1867–1882, 2016.

[6] Stefan Peidli, Tessa D Green, Ciyue Shen, Torsten Gross, Joseph Min, Samuele Garda, Bo Yuan, Linus J Schumacher, Jake P Taylor-King, Debora S Marks, et al. scperturb: harmonized single-cell perturbation data. Nature Methods, 21(3):531–540, 2024.

[7] Sanjay R Srivatsan, José L McFaline-Figueroa, Vijay Ramani, Lauren Saunders, Junyue Cao, Jonathan Packer, Hannah A Pliner, Dana L Jackson, Riza M Daza, Lena Christiansen, et al. Massively multiplex chemical transcriptomics at single-cell resolution. Science, 367(6473):45–51, 2020.

[8] Jesse Zhang, Airol A Ubas, Richard de Borja, Valentine Svensson, Nicole Thomas, Neha Thakar, Ian Lai, Aidan Winters, Umair Khan, Matthew G Jones, et al. Tahoe-100m: A giga-scale single-cell perturbation atlas for context-dependent gene function and cellular modeling. BioRxiv, pages 2025–02, 2025.

[9] Ajay Nadig, Joseph M. Replogle, Angela N. Pogson, Mukundh Murthy, Steven A. McCarroll, Jonathan S. Weissman, Elise B. Robinson, and Luke J. O’Connor. Transcriptome-wide analysis of differential expression in perturbation atlases. Nature Genetics, 57(5):1228–1237, May 2025.

[10] Artur Szałata, Andrew Benz, Robrecht Cannoodt, Mauricio Cortes, Jason Fong, Sunil Kuppasani, Richard Lieberman, Tianyu Liu, Javier A. Mas-Rosario, Rico Meinl, Jalil Nourisa, Jared Tumiel, Tin M. Tunjic, Mengbo Wang, Noah Weber, Hongyu Zhao, Benedict Anchang, Fabian J. Theis, Malte D. Luecken, and Daniel B. Burkhardt. A benchmark for prediction of transcriptomic responses to chemical perturbations across cell types. Advances in Neural Information Processing Systems, 37:20566–20616, December 2024.

[11] Benjamin DeMeo, Charlotte Nesbitt, Samuel A. Miller, Daniel B. Burkhardt, Inna Lipchina, Doris Fu, Peter Holderrieth, David Kim, Sergey Kolchenko, Artur Szalata, Ishan Gupta, Christine Kerr, Thomas Pfefer, Raziel Rojas-Rodriguez, Sunil Kuppassani, Laurens Kruidenier, Parul B. Doshi, Mahdi Zamanighomi, James J. Collins, Alex K. Shalek, Fabian J. Theis, and Mauricio Cortes. Active learning framework leveraging transcriptomics identifies modulators of disease phenotypes. Science, 390(6776):eadi8577, October 2025.

[12] Mitchell L. Leibowitz, Stamatis Papathanasiou, Phillip A. Doerfler, Logan J. Blaine, Lili Sun, Yu Yao, Cheng-Zhong Zhang, Mitchell J. Weiss, and David Pellman. Chromothripsis as an ontarget consequence of CRISPR–Cas9 genome editing. Nature Genetics, 53(6):895–905, June 2021.

[13] Longda Jiang, Carol Dalgarno, Efthymia Papalexi, Isabella Mascio, Hans-Hermann Wessels, Huiyoung Yun, Nika Iremadze, Gila Lithwick-Yanai, Doron Lipson, and Rahul Satija. Systematic reconstruction of molecular pathway signatures using scalable single-cell perturbation screens. Nature Cell Biology, 27(3):505–517, March 2025.

[14] Po-Yuan Tung, John D. Blischak, Chiaowen Joyce Hsiao, David A. Knowles, Jonathan E. Burnett, Jonathan K. Pritchard, and Yoav Gilad. Batch effects and the effective design of single-cell gene expression studies. Scientific Reports, 7(1):39921, January 2017.

[15] Wenxin Long, Tianyu Liu, Lingzhou Xue, and Hongyu Zhao. spvelo: Rna velocity inference for multi-batch spatial transcriptomics data. Genome Biology, 26(1):239, 2025.

[16] Mark-Anthony Bray, Shantanu Singh, Han Han, Chadwick T Davis, Blake Borgeson, Cathy Hartland, Maria Kost-Alimova, Sigrun M Gustafsdottir, Christopher C Gibson, and Anne E Carpenter. Cell painting, a high-content image-based assay for morphological profiling using multiplexed fluorescent dyes. Nature Protocols, 11(9):1757–1774, 2016.

[17] Srinivas Niranj Chandrasekaran, Beth A Cimini, Amy Goodale, Lisa Miller, Maria Kost-Alimova, Nasim Jamali, John G Doench, Briana Fritchman, Adam Skepner, Michelle Melanson, et al. Three million images and morphological profiles of cells treated with matched chemical and genetic perturbations. Nature Methods, 21(6):1114–1121, 2024.

[18] Marta M Fay, Oren Kraus, Mason Victors, Lakshmanan Arumugam, Kamal Vuggumudi, John Urbanik, Kyle Hansen, Safiye Celik, Nico Cernek, Ganesh Jagannathan, et al. Rxrx3: Phenomics map of biology. BioRxiv, pages 2023–02, 2023.

[19] Leon Hetzel, Simon Boehm, Niki Kilbertus, Stephan Günnemann, Fabian Theis, et al. Predicting cellular responses to novel drug perturbations at a single-cell resolution. Advances in Neural Information Processing Systems, 35:26711–26722, 2022.

[20] Yusuf Roohani, Kexin Huang, and Jure Leskovec. Predicting transcriptional outcomes of novel multigene perturbations with gears. Nature Biotechnology, 42(6):927–935, 2024.

[21] Zoe Piran, Niv Cohen, Yedid Hoshen, and Mor Nitzan. Disentanglement of single-cell data with biolord. Nature Biotechnology, 42(11):1678–1683, 2024.

[22] Gefei Wang, Tianyu Liu, Jia Zhao, Youshu Cheng, and Hongyu Zhao. Modeling and predicting single-cell multi-gene perturbation responses with sclambda. BioRxiv, 2024.

[23] Dominik Klein, Jonas Simon Fleck, Daniil Bobrovskiy, Lea Zimmermann, Sören Becker, Alessandro Palma, Leander Dony, Alejandro Tejada-Lapuerta, Guillaume Huguet, Hsiu-Chuan Lin, et al. Cellflow enables generative single-cell phenotype modeling with flow matching. BioRxiv, pages 2025–04, 2025.

[24] Abhinav K Adduri, Dhruv Gautam, Beatrice Bevilacqua, Alishba Imran, Rohan Shah, Mohsen Naghipourfar, Noam Teyssier, Rajesh Ilango, Sanjay Nagaraj, Mingze Dong, et al. Predicting cellular responses to perturbation across diverse contexts with state. BioRxiv, pages 2025–06, 2025.

[25] Tianyu Liu, Tianqi Chen, Wangjie Zheng, Xiao Luo, Yiqun Chen, and Hongyu Zhao. Embeddings from language models are good learners for single-cell data analysis. Patterns, page 101431, 2026.

[26] Yuchen Wang, Tianchi Lu, Xingjian Chen, Zhongyu Yao, and Ka-Chun Wong. scotm: a deep learning framework for predicting single-cell perturbation responses with large language models. Bioengineering, 12(8):884, 2025.

[27] Haotian Cui, Chloe Wang, Hassaan Maan, Kuan Pang, Fengning Luo, Nan Duan, and Bo Wang. scgpt: toward building a foundation model for single-cell multi-omics using generative ai. Nature Methods, 21(8):1470–1480, 2024.

[28] Minsheng Hao, Jing Gong, Xin Zeng, Chiming Liu, Yucheng Guo, Xingyi Cheng, Taifeng Wang, Jianzhu Ma, Xuegong Zhang, and Le Song. Large-scale foundation model on single-cell transcriptomics. Nature Methods, 21(8):1481–1491, 2024.

[29] Christina V Theodoris, Ling Xiao, Anant Chopra, Mark D Chaffin, Zeina R Al Sayed, Matthew C Hill, Helene Mantineo, Elizabeth M Brydon, Zexian Zeng, X Shirley Liu, et al. Transfer learning enables predictions in network biology. Nature, 618(7965):616–624, 2023.

[30] Artur Szałata, Karin Hrovatin, Sören Becker, Alejandro Tejada-Lapuerta, Haotian Cui, Bo Wang, and Fabian J. Theis. Transformers in single-cell omics: a review and new perspectives. Nature Methods, 21(8):1430–1443, August 2024.

[31] Tianyu Liu, Kexing Li, Yuge Wang, Hongyu Li, and Hongyu Zhao. Evaluating the utilities of foundation models in single-cell data analysis. BioRxiv, pages 2023–09, 2023.

[32] Kasia Z Kedzierska, Lorin Crawford, Ava P Amini, and Alex X Lu. Zero-shot evaluation reveals limitations of single-cell foundation models. Genome Biology, 26(1):101, 2025.

[33] Mohammad Lotfollahi, F Alexander Wolf, and Fabian J Theis. scgen predicts single-cell perturbation responses. Nature Methods, 16(8):715–721, 2019.

[34] Hanwen Xing and Christopher Yau. Gperturb: Gaussian process modelling of single-cell perturbation data. Nature Communications, 16(1):5423, 2025.

[35] Conrad L Schoch, Stacy Ciufo, Mikhail Domrachev, Carol L Hotton, Sivakumar Kannan, Rogneda Khovanskaya, Detlef Leipe, Richard Mcveigh, Kathleen O’Neill, Barbara Robbertse, et al. Ncbi taxonomy: a comprehensive update on curation, resources and tools. Database, 2020:baaa062, 2020.

[36] Hao Chen, Frederick J King, Bin Zhou, Yu Wang, Carter J Canedy, Joel Hayashi, Yang Zhong, Max W Chang, Lars Pache, Julian L Wong, et al. Drug target prediction through deep learning functional representation of gene signatures. Nature Communications, 15(1):1853, 2024.

[37] TahoeBlog. Target deconvolution through data integration: Unifying drug and genetic perturbations. https://blog.tahoebio.ai/p/target-deconvolution-through-data. Accessed: 2026-01-21.

[38] David DeTomaso, Matthew G Jones, Meena Subramaniam, Tal Ashuach, Chun J Ye, and Nir Yosef. Functional interpretation of single cell similarity maps. Nature Communications, 10(1):4376, 2019.

[39] Kaspar Märtens, Marc Boubnovski Martell, Cesar A Prada-Medina, and Rory Donovan-Maiye. Langpert: Llm-driven contextual synthesis for unseen perturbation prediction. In ICLR 2025 Workshop on Machine Learning for Genomics Explorations.

[40] Joseph M Replogle, Reuben A Saunders, Angela N Pogson, Jeffrey A Hussmann, Alexander Lenail, Alina Guna, Lauren Mascibroda, Eric J Wagner, Karen Adelman, Gila Lithwick-Yanai, et al. Mapping information-rich genotype-phenotype landscapes with genome-scale perturb-seq. Cell, 185(14):2559–2575, 2022.

[41] Yiqun Chen and James Zou. Genepert: Leveraging genept embeddings for gene perturbation prediction. BioRxiv, pages 2024–10, 2024.

[42] Elijah Cole, Geert-Jan Huizing, Sohan Addagudi, Nicholas Ho, Euxhen Hasanaj, Merel Kuijs, Toby Johnstone, Maria Carilli, Alec Davi, Caleb Ellington, et al. Foundation models improve perturbation response prediction. BioRxiv, pages 2026–02, 2026.

[43] Yiqun Chen and James Zou. Simple and effective embedding model for single-cell biology built from chatgpt. Nature Biomedical Engineering, 9(4):483–493, 2025.

[44] Damian Szklarczyk, Rebecca Kirsch, Mikaela Koutrouli, Katerina Nastou, Farrokh Mehryary, Radja Hachilif, Annika L Gable, Tao Fang, Nadezhda T Doncheva, Sampo Pyysalo, et al. The string database in 2023: protein–protein association networks and functional enrichment analyses for any sequenced genome of interest. Nucleic Acids Research, 51(D1):D638–D646, 2023.

[45] Ann C Huang, Tsung-Han S Hsieh, Jiang Zhu, Jackson Michuda, Ashton Teng, Soohong Kim, Elizabeth M Rumsey, Sharon K Lam, Ikenna Anigbogu, Philip Wright, et al. X-atlas/orion: Genome-wide perturb-seq datasets via a scalable fix-cryopreserve platform for training dose-dependent biological foundation models. BioRxiv, pages 2025–06, 2025.

[46] Tianyu Liu, Edward De Brouwer, Tony Kuo, Nathaniel Diamant, Alsu Missarova, Hanchen Wang, Minsheng Hao, Hector Corrada Bravo, Gabriele Scalia, Aviv Regev, et al. Learning multi-cellular representations of single-cell transcriptomics data enables characterization of patient-level disease states. In International Conference on Research in Computational Molecular Biology, pages 303–306, 2025.

[47] Mingyu Lu, Ethan Weinberger, Chanwoo Kim, and Su-In Lee. Cellclip-learning perturbation effects in cell painting via text-guided contrastive learning. Advances in Neural Information Processing Systems, 38:124505–124537, 2026.

[48] Oren Kraus, Federico Comitani, John Urbanik, Kian Kenyon-Dean, Lakshmanan Arumugam, Saber Saberian, Cas Wognum, Safiye Celik, and Imran S Haque. Rxrx3-core: Benchmarking drug-target interactions in high-content microscopy. arXiv preprint arXiv:2503.20158, 2025.

[49] Maxime Oquab, Timothée Darcet, Théo Moutakanni, Huy Vo, Marc Szafraniec, Vasil Khalidov, Pierre Fernandez, Daniel Haziza, Francisco Massa, Alaaeldin El-Nouby, et al. Dinov2: Learning robust visual features without supervision. arXiv preprint arXiv:2304.07193, 2023.

[50] F Alexander Wolf, Philipp Angerer, and Fabian J Theis. Scanpy: large-scale single-cell gene expression data analysis. Genome biology, 19(1):15, 2018.

[51] MedChemExpress. Medchemexpress. Accessed: 2026-02-04.

[52] Selenium Contributers. The selenium project. Accessed: 2026-02-04.

[53] Alec Radford, Jong Wook Kim, Chris Hallacy, Aditya Ramesh, Gabriel Goh, Sandhini Agarwal, Girish Sastry, Amanda Askell, Pamela Mishkin, Jack Clark, et al. Learning transferable visual models from natural language supervision. In International Conference on Machine Learning, pages 8748–8763. PMLR, 2021.

[54] Xiaohua Zhai, Basil Mustafa, Alexander Kolesnikov, and Lucas Beyer. Sigmoid loss for language image pre-training. In Proceedings of the IEEE/CVF International Conference on Computer Vision, pages 11975–11986, 2023.

[55] Aravind Subramanian, Pablo Tamayo, Vamsi K Mootha, Sayan Mukherjee, Benjamin L Ebert, Michael A Gillette, Amanda Paulovich, Scott L Pomeroy, Todd R Golub, Eric S Lander, et al. Gene set enrichment analysis: a knowledge-based approach for interpreting genome-wide expression profiles. Proceedings of the National Academy of Sciences, 102(43):15545–15550, 2005.

[56] Arthur Liberzon, Chet Birger, Helga Thorvaldsdóttir, Mahmoud Ghandi, Jill P Mesirov, and Pablo Tamayo. The molecular signatures database hallmark gene set collection. Cell Systems, 1(6):417–425, 2015.

[57] Matthew Cannon, James Stevenson, Kathryn Stahl, Rohit Basu, Adam Coffman, Susanna Ki-wala, Joshua F McMichael, Kori Kuzma, Dorian Morrissey, Kelsy Cotto, et al. Dgidb 5.0: rebuilding the drug–gene interaction database for precision medicine and drug discovery platforms. Nucleic Acids Research, 52(D1):D1227–D1235, 2024.

